# Surveying Brain Tumor Heterogeneity by Single-Cell RNA Sequencing of Multi-sector Biopsies

**DOI:** 10.1101/2020.01.19.911701

**Authors:** Kai Yu, Yuqiong Hu, Fan Wu, Qiufang Guo, Zenghui Qian, Waner Hu, Jing Chen, Kuanyu Wang, Xiaoying Fan, Xinglong Wu, John EJ Rasko, Xiaolong Fan, Antonio Iavarone, Tao Jiang, Fuchou Tang, Xiao-Dong Su

## Abstract

Brain tumors are among the most challenging human tumors for which the mechanisms driving progression and heterogeneity remain poorly understood. We combined single-cell RNA-seq with multisector biopsies to sample and analyze single-cell expression profiles of gliomas from 13 Chinese patients. After classifying individual cells, we generated a spatial and temporal landscape of glioma that revealed the patterns of invasion between the different sub-regions of gliomas. We also used single-cell inferred CNVs and pseudotime trajectories to inform on the crucial branches that dominate tumor progression. The dynamic cell components of the multi-region biopsy analysis allowed us to spatially deconvolute with unprecedented accuracy the transcriptomic features of the core and those of the periphery of glioma at single cell level. Through this rich and geographically detailed dataset, we were also able to characterize and construct the chemokine and chemokine receptor interactions that exist among different tumor and non-tumor cells. This study provides the first spatial-level analysis of the cellular states that characterize human gliomas. It also presents an initial molecular map of the crosstalks between glioma cells and the surrounding microenvironment with single cell resolution.

## INTRODUCTION

Gliomas are among the most lethal forms of human tumors as they are characterized by aggressive behaviors and resistance to multiple therapies. The development of genetic mutations in malignant cells and the complex interactions between tumor and non-tumor cells in the glioma micro-environment foster intratumoral heterogeneity, thus contributing to the therapeutic failures and the generally poor prognosis of gliomas. A major unmet challenge in neuro-oncology is our ability to understand glioma heterogeneity and progression in gliomas and how they influence therapeutic resistance [1].

Several studies reported that malignant gliomas are characterized by a formidable degree of intratumoral heterogeneity. For example, mosaic amplification of receptor tyrosine kinase genes *(EGFR, MET, PDGFRA)* is known to represent a classical hallmark of genetic heterogeneity affecting neighboring tumor cells within bulk glioma samples [2]. Furthermore, single cell-derived clones of glioma cells have been identified and shown to exhibit divergent proliferation and differentiation abilities [3]. Finally, the multi-region genetic analysis of gliomas with single nucleotide polymorphism (SNP) array or whole exome sequencing revealed that divergent glioma subtypes can be recovered from different geographical regions, which together give rise to a branched pattern of progression [4,5]. As single-cell RNA-sequencing became a feasible approach to investigate human tumors, glioma heterogeneity has started to be explored with single-cell resolution [6–11]. However, most of the studies that have previously reported single cell RNA-sequencing of gliomas did not include the analysis of either tumor cells and the tumor microenvironment (TME) from multiple spatially annotated regions of gliomas, thus limiting our understanding of patterns of spatial evolution and brain infiltration, the latter being one the most critical hallmark of aggressiveness and progression of malignant glioma.

To delineate the glioma single-cell heterogeneity in both spatial and temporal resolution, we performed a glioma single-cell analysis from multisector biopsies informed by precision navigation surgery. Cell type components of each tumor fragments and temporal relationship of cells in each individual patient were unbiasedly identified. Our analysis did not use approaches aimed at selecting specific tumor or non-tumor cell populations. Therefore, we report the first single cell based comprehensive spatial analysis of the geographical structure of glioma and the dynamic progression of the interactions of tumor cells with individual non-tumor cells from multiple tumor locations.

## RESULTS

### Precision Navigation Based Multi-sector Biopsies and Single-Cell RNA-seq of Glioma Cells

Tumor sections with potential representative divergent properties were marked in a presurgical 3D enhanced magnetic resonance imaging (MRI) model and tumor tissues were precisely collected during surgery by navigation sampling. Samples were quickly dissociated and subjected to single-cell RNA sequencing (scRNA-seq) library preparation [12,13] (Fig. 1A). Overall, 7,928 single-cell transcriptomes were generated, and 6,148 passed stringent quality filtering steps after alignment and reads counting (Fig. 1B and Supplementary Fig. S1A-B). These cells were collected from 73 regions in 13 patients with glioma covering the most frequent subtypes (3 WHO grade II, 1 WHO grade III, 8 WHO grade IV and 1 gliosarcoma). As control, we also included one brain metastasis from a patient with lung squamous cell carcinoma (Fig. 1C, Supplementary Fig. S2, Supplementary Table 1 and 2).

**Figure 1.**
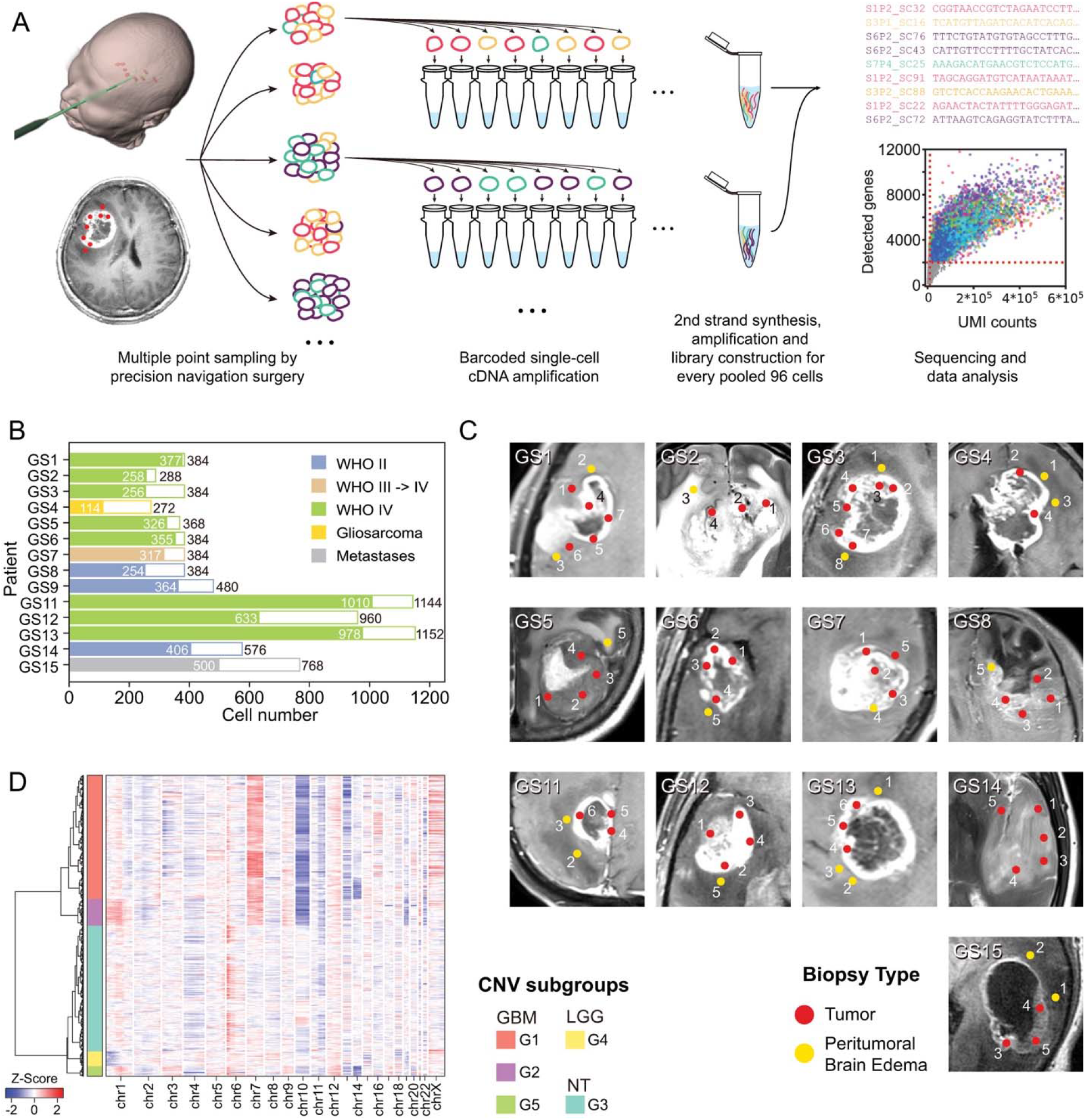
General information of experiment procedure and generated data. (A) Experiment procedure. Multiple point sampling was done by precision navigation surgery, followed by single-cell isolation and barcoded single-cell cDNA amplification. After every 96 cells were pooled together, the sequencing library was constructed by several experiment procedures. (B) Pathology information and cell number of 14 patients collected in this study. White bars and colored bars represented raw cell number and filtered cell number in each patient, respectively. (C) Medical imaging and biopsy locations of all sampling points in each patient. Red and yellow dots marked locations of tumoral and peritumoral sampling point in the MRI image. (D) RNA-derived single cell CNVs information. Hierarchical clustering divided all glioma related cells into 5 CNV subtypes.

Since regional gene expression status could be affected by copy number variation (CNV) dose effect, we adopted previously reported methods to predict large fragment copy number status with single-cell gene expression matrix [6,8]. The generated copy number matrix clustered into 5 CNV subgroups (Fig. 1D). Malignant cells were identified based on this classification. Subgroup G1 and G2 included glioblastoma multiforme (GBM) cells sharing a *chr7^amp^/chr10^del^* driven transcriptome whereas a unique CNV pattern was apparent in GBM patient GS3 (subgroup G5). The G4 subgroup included low-grade glioma (LGG) cells with *chr1p/chr19q* codeletion-driven signatures. The last subgroup (G3) was composed of non-malignant cells without obvious CNVs.

### Optimized t-SNE Map and Clustering Identifies 25 Cell Type Clusters

As reported by previous single cell RNA-seq studies of human glioma, the malignant cells from different patients showed a fragmented relationship in clustering analysis, mainly because of the dose effect of gene expression caused by diverse CNVs [6,8]. Therefore, we argued that removal of CNV variances would optimize the principal component analysis (PCA) and t-Distributed Stochastic Neighbor Embedding (t-SNE) analysis.

To obtain the CNV status of each tumor sample, we performed low depth whole genome sequencing (WGS) on bulk biopsies. Based on the WGS data, genes with interpatient large copy number changes were removed from the clustering analysis (Fig. 2A, Supplementary Fig. S3B and S4). The integrated analysis of tumor cells from our entire dataset using PCA and t-SNE resulted in an optimized map of 25 cell clusters (Fig. 2C,D and Supplementary Fig. S5). Compared to the original t-SNE map (Fig. 2B and Supplementary Fig. S6A, B and D), the fragmented cell distribution was resolved to a map with more concentrated points and clearer edges (Fig. 2D). This distribution was confirmed by independent binary regulon activity clustering with SCENIC (Fig. 2E and Supplementary Fig. S6E and S7) [14]. Combination of these different strategies could help minimize analyze artifact in analysis.

**Figure 2.**
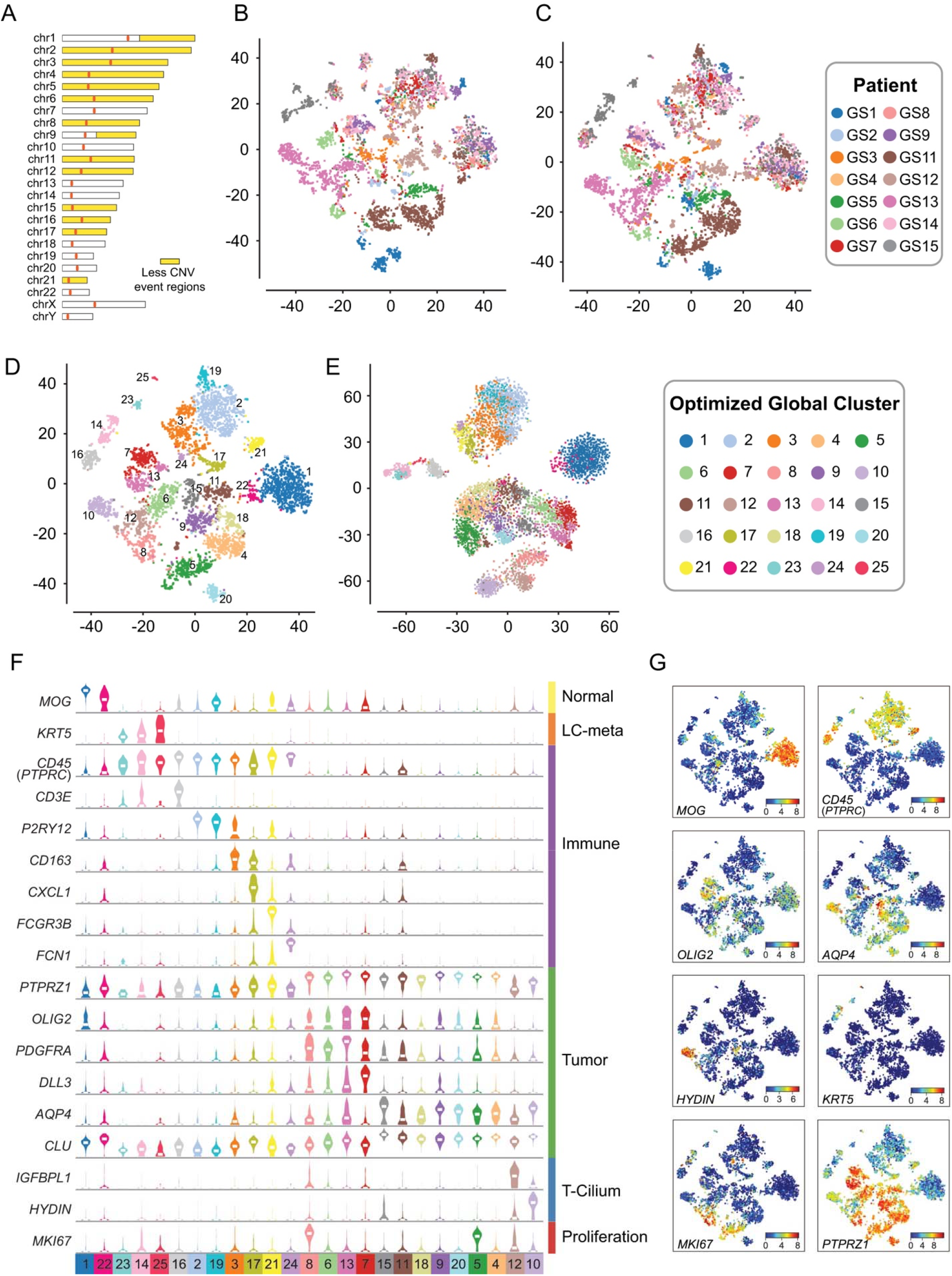
Molecular characteristics of cells from glioma. (A) Based on bulk CNVs status, chromosome regions that had less CNVs events (1.8<copy<2.2, abnormal patient<3) in our samples were marked by yellow color, and variable genes in these regions were used for PCA and t-SNE analysis. (B) Pre-optimize global t-SNE map, colored by patients. (C) After-optimize global t-SNE map, colored by patients. (D) Global map of sub-clusters using t-SNE coordinates. Dots were individual cells and colored by optimized global clusters. (E) t-SNE analysis based on binary regulon activity, analyzed by SCENIC. Clusters had similar distribution with global 25 cluster t-SNE map. (F) was according to (D). With shared and specifically expressed patterns showed in (G), sub-clusters could be divided into normal *(MOG^+^),* immune (*CD45*^+^), regular glioma tumor *(PTPRZ1^+^, OLIG2^+^* or *AQP4^+^*), cilia property tumor *(HYDIN^+^),* proliferation tumor *(MKI67^+^)* and metastasis cells.

Considering the cell source and CNV status with a global t-SNE map, the non-tumor cells and the glioma cells lacking extensive genomic rearrangements (such as those derived from LGG) showed a converged distribution, indicating that our data were not influenced by batch effects. They also indicated that the LGG-derived tumor cells were more uniform among different patients (Fig. 2C and Supplementary Fig. S6C). Cell cluster-specific genes coalesced the 25 clusters into 4 major groups (Supplementary Table 2): *CNV^-^MOG^+^* normal glial cells [15], *KRT5^+^* lung cancer (LC) metastasis cells [16], *CD45 (PTPRC)*^+^ immune cells and *CNV^+^* malignant tumor cells. In malignant glioma cells, *OLIG2/DLL3* and *AQP4/CLU* distinguish tumor cells exhibiting transcriptomic features of oligodendrocytes and astrocytes, respectively [17–19]. Cells positive to the general proliferative marker *MKI67*^+^ were present among both glioma cell phenotypes. Moreover, *PTPRZ1* and *SOX2* were significantly overexpressed in glioma cells and could therefore be considered as useful markers to estimate the tumor cell purity in bulk glioma tissues. Interestingly, clusters 10 and 12 specifically overexpressed cilium-related markers *(HYDIN, FOXJ1,* etc.) [20,21]. The tumor cells included in these clusters exhibited astrocyte-like features and expressed low levels of *PTPRZ1* (Fig. 2F, G and Supplementary Fig. S8). Clusters 10 and 12 mostly derived from patient GS13, indicating that this patient developed a rare form of ciliated glioma for which single-cell studies have not previously been reported.

### Annotation of the Glioma Microenvironment and TCGA-based Classification of Single Glioma Cells

Normal diploid cells in the tumor samples were mainly cells within the tumor microenvironment (TME). To conduct an unbiased investigation of the TME in glioma, we did not include flow cytometry-based selection in our sampling procedure. Normal oligodendrocytes, microglia and macrophages accounted for approximately half of these cells, with relative homogeneity among patients, such a conclusion could not have emerged from *CD45* selected samples, as this is the prevailing sampling method used in previous scRNA-seq studies. We also captured cells showing specialized activities, such as interferon-induced oligodendrocytes and polarized microglia/macrophages (Fig. 3A).

**Figure 3.**
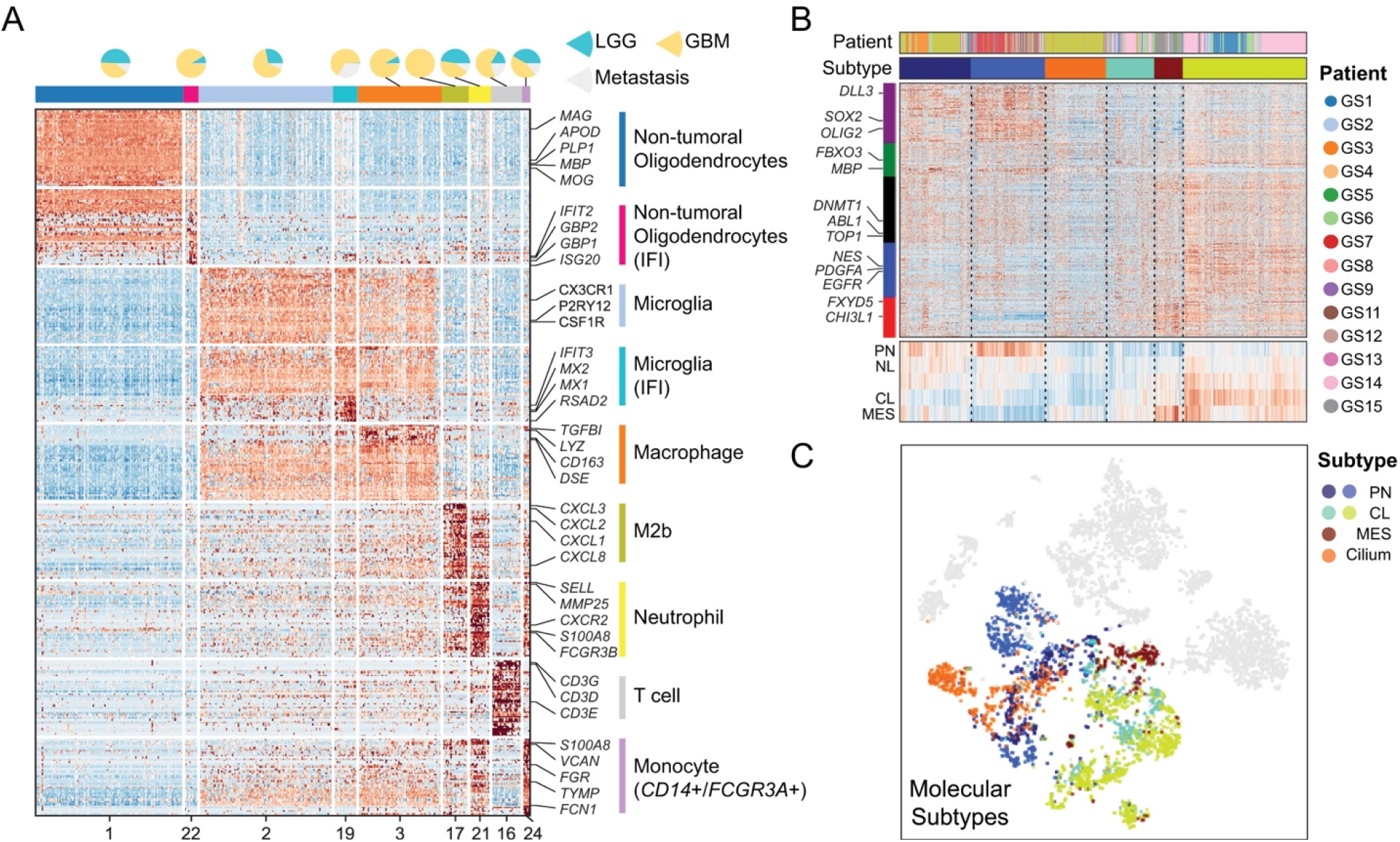
Cell type annotation of tumor microenvironment cells and malignant cells. (A) Heatmap showed marker genes of sub-clusters which were tumor microenvironment cells. Pie plot at the top showed cell ratio of LGG/GBM/Metastasis samples. Cells in clusters marked with IFI showed interferon induced property. (B) Cell clustered by TCGA 4 classification genes. Heatmap at the bottom was the average expression value of 5 gene-sets marked in the left color bar. (C) Tumor cell subtype mapped in t-SNE coordinates.

Next, we classified malignant cells according to the The Cancer Genome Atlas (TCGA) GBM classification model [19]. Grade II tumor cells with chromosome *1p/19q* codeletion had a strong proneural (PN) signature. Conversely, the individual tumor cells recovered from GBM were distributed among each of the three transcriptomic subgroups (PN, classical (CL) or mesenchymal (MES)). We did not identify glioma cells that could be assigned to the neural (NL) subtype but we detected cells expressing cilium-related signatures that were distinct from those associated with known TCGA subgroups (Fig. 3B and C).

### CNV Accumulation and Tumor Progression in Patient GS1

Since CNVs usually accumulate during tumor progression, the CNV profile of individual tumor cells has emerged as an accurate inference to trace tumor progression [2–4,8,22,23]. We used arm-level scRNA-seq derived CNVs to trace tumor cell clones, which were confirmed by bulk WGS (Fig. 1D, Supplementary Fig. S3A and B) to avoid possible mistakes by unravel single-cell CNV from RNA data with bioinformatics tools alone. Multiple cell subpopulations with accumulated CNVs were found in patients GS1, GS13 and GS5 (Supplementary Fig. S3C and S9).

From patient GS1, a female with an *IDH-wild* type GBM, single cells were collected from 5 tumor core locations (P1/4/5/6/7) and 2 peritumoral locations (P2/3). A few cells (P8) were also collected from an adjacent brain region that from the imaging analysis lacked evidence of tumor infiltration. Ring plots illustrate the tumor and non-tumor cell components at each site (Fig. 4A). In this patient, CL glioma cells accounted for most of the malignant component at all tumor sites, with a small fraction of PN tumor cells. Interestingly, we found that macrophages composed the larger fraction of non-tumor cells infiltrating the tumor core (P1/4/5/6/7) but they were replaced by microglia in the peritumoral invading front (P2/3). The peritumoral biopsies contained a higher fraction of non-tumor cells than glioma cells.

**Figure 4.**
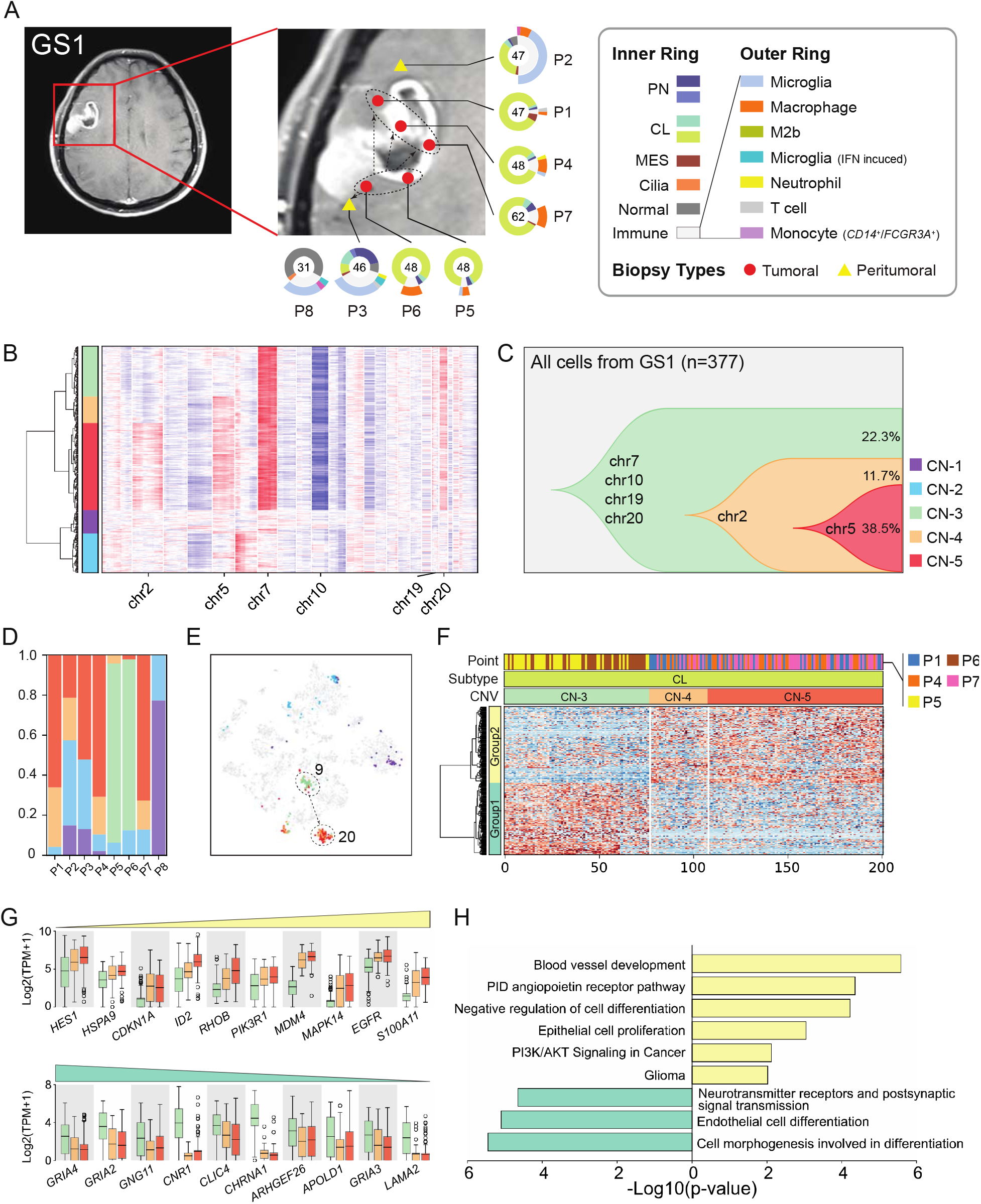
Glioma clonal evolution of Patient GS1 at spatial and temporal resolution. (A) MRI image of Patient GS1. Yellow and red markers in zoomed image represented peritumoral and tumoral sampling points. Ring plot in the right and bottom displayed cell components of each point. Color of the inner ring showed classified glial cell subtypes, and the outer ring showed detailed immune cell subtypes. Cell numbers were labeled in the center of these ring plots. (B) Single-cell CNV heatmap. Cells were divided into 5 groups by hierarchical clustering. (C) Clonal evolution trail followed by accumulating CNV events. Each color represented a CNV subclone and chromosomes were labeled, in which copy number alteration were happened during clonal transition. (D) CNV subclone components in each sampling point, P5/6 had a different component compared with other tumoral/peritumoral points. (E) CNV subclone distributions in t-SNE coordinates. When CN-3 developed into CN-4/5, cells translocated from cluster 9 to cluster 20. (F) Heatmap of differentially expressed genes between CN-3/4/5 CL glial cells. Differential genes could be divide into 2 gene sets, marked with the left color bar. (G) Box plot of differential genes. The increased (upper) and decreased (lower) gene expression profiles followed CNV accumulation. Boxes were colored by CNV subgroup. (H) Functional enrichment of increased and decreased gene-sets.

As scRNA-seq can be used to infer the underlying CNV, we interrogated our dataset to uncover the spatial dynamics of CNV changes in glioma cells in order to reconstruct the trajectory of tumor initiation and progression. Five CNV-based clusters (CN-1 to CN-5) were found in patient GS1 (Fig. 4B). CN-1 and CN-2 contained diploid non-tumor cells, but were divided into two clusters because their expression profiles differed from those of normal glial and immune cells. CN-3 to CN-5 clusters contained a series of aneuploid subclones that shared the *chr7/10/19/20* CNVs that are recurrent genomic alterations in GBM. As they progressed, these subclones first accumulated CNVs on *chr2* and later on *chr5* as they transitioned from CN-3 to CN-4 and from CN-4 to CN-5, respectively (Fig. 4C), therefore highlighting the temporal progression of this tumor. The combination of spatial and temporal information indicated that the CN-3 population was distributed only in P5/P6, whereas the CN-4 and CN-5 populations were present in all other tumoral and peritumoral sites (Fig. 4D). These findings clearly indicate that P5/P6 were the initial sites of the tumor that progressed towards the locations of P1/P4/P7 and P3 (Fig. 4A). In the global t-SNE analysis, as clone CN-3 developed into CN-4 and CN-5, glioma cells also transformed from cluster 9 to 20 (Fig. 4E). Our findings established that the number of somatic CNVs increase across the different glioma regions, thus defining a pattern of tumor progression. They are also consistent with the notion that an increased definition of the multi-sector sampling of glioma might reveal finer and more accurate genetic trajectories of glioma evolution, as shown in recent hepato-carcinoma studies [24].

To unravel the phenotypic changes that mark spatial evolution of glioma we performed differential gene expression analysis between the different clones. As cells classified within the CN-3 clone were replaced by those in the CN-4 and CN-5 clones, we observed increased expression of genes implicated in negative regulation of cell differentiation *(HES1, HSPA9, ID2),* DNA damage response *(MDM4, SOX4)* and chemoattractant cytokine and neutrophil activation *(MAPK14, S100A11),* by general function annotation. The transition from CN-3 to CN-4 and CN-5 clones coincided also with increased expression of the *EGFR* oncogene. Conversely, genes expressed in benign astrocytes *(CNR1, CHRNA1, LAMA2, GNG11, GRIA2, GRIA3, GRIA4)* and general endothelial cell differentiation markers *(ARHGEF26, CLIC4, APOLD1)* were suppressed during the transition (Fig. 4F and G, Supplementary Table 2). Considering the functional enrichment of differentially expressed genes, our findings converge on a model whereby the genetic alterations such as CNVs that accumulated during turn or invasion lead to loss of differentiated astrocyte properties and gain of known features driving tumor aggressiveness, angiogenesis, dedifferentiation, oncogenic *PI3K/AKT* signaling (Fig. 4H), ultimately resulting in the promotion of glioma progression towards more aggressive and invasive phenotypes.

### The Trajectory of Tumor Cell States Reveals Branched Progression in Patient GS13

Patient GS13 was a male with an *IDH-wild* type GBM characterized by high expression of genes associated with motile cilium activities (e.g., *FOXJ1, FAM183A*, *HYDIN, DNALI1,* etc.) (Supplementary Fig. S8 and S10). We noted that only about 5% of TCGA GBM patients exhibited high expression of cilium-related genes. Therefore, the particular type of GBM analyzed from patient GS13 belongs to a rare type of glioma that was not previously investigated at single cell level. However, cilium-specific gene expression was not associated with a specific pattern of survival (Supplementary Fig. S11). From this patient, we acquired 978 cells from 3 core and 2 peritumoral sites, and found that the cells at core locations consisted of PN and cilium-positive cells (Fig. 5A). CNVs were also evident in this case. The clonal CNV reconstruction revealed 2 CNV clones (CN-2 and CN-3) and the transition from CN-2 to CN-3 was marked by accumulation of *chr1^amp^* and *chr19^del^.* Furthermore, as cells transitioned to CN-3, the expression profile changed from global cluster 6 (PN) to global clusters 10 and 12 (cilium) (Fig. 5B-E). We also applied hierarchical clustering of CN-2 and CN-3 malignant cells and identified differentially expressed genes between these two groups. Interestingly, after cells transitioned to CN-3, they formed 2 branches with distinct gene expression profiles (Fig. 5F and H, Supplementary Table 2). The functional gene set enrichment analysis showed that the transition from CN-2 to CN-3 was associated with decreased expression of group 1 genes, which were implicated in glial cell differentiation and adhesion. Conversely, the expression of genes in groups 2 and 3 (implicated in cell cycle/DNA replication and cilium regulation, respectively) increased (Fig. 5G and Supplementary Fig. S12).

**Figure 5.**
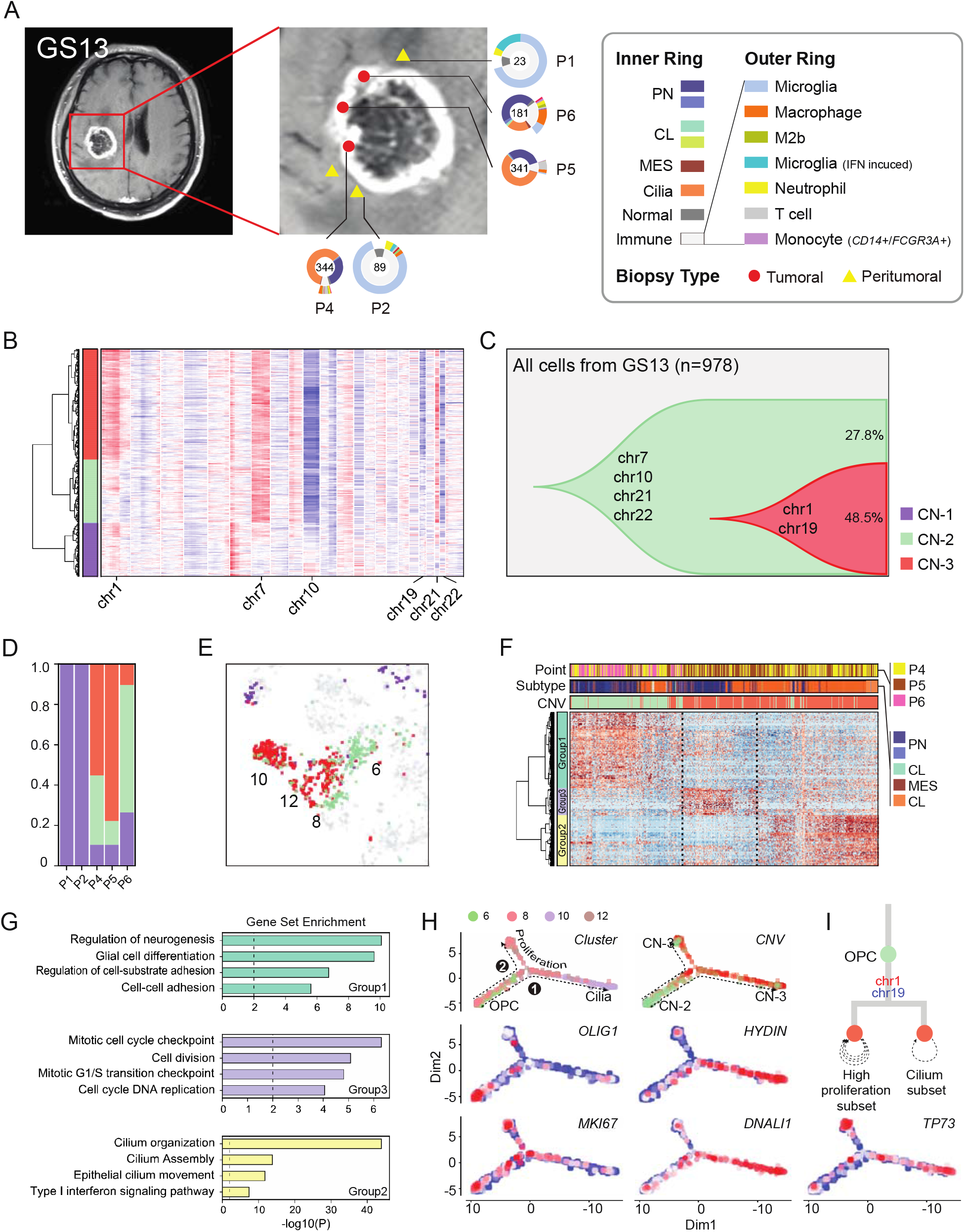
Glioma clonal evolution of Patient GS13 at spatial and temporal resolution. (A) MRI image of Patient GS13. Yellow and red markers in zoomed image represented peritumoral and tumoral sampling points. Ring plot in the right and bottom displayed cell components of each point. Color of the inner ring showed classified glial cell subtypes, and the outer ring showed detailed immune cell subtypes. Cell numbers were labeled in the center of these ring plots. (B) Single-cell CNV heatmap, cells were divided into 3 groups by hierarchical clustering. (C) Clonal evolution trail followed by accumulating CNV events. Each color represented a CNV subclone and chromosomes were labeled, in which copy number alterations happened during clonal transition. (D) CNV subclones components in each sampling point, P6 had a lower CN-3 ratio than P4/5. (E) CNV subclones distribution in t-SNE coordinates. When CN-2 developed into CN-3, cells translocated from cluster 6 to cluster 12/10. (F) Heatmap of differentially expressed genes between CN-2/5 glial cells. Hierarchical clustering were applied in both gene and cell dimension. Differential genes could be divide into 3 gene sets, marked with the left color bar. (G) Functional enrichment of 3 gene-sets in (F). (H) Single-cell trajectories of malignant cells in GS13. Top left subplot was colored by global t-SNE clusters, 2 branches of cells were developed from OPC cells. The top right subplot was colored by CNV groups. The remaining 4 subplots were relative expression patterns of marker genes *(OLIG2, HYDIN, MKI67, DNALI1* and *TP73).* (I) Branched clonal developing model of GS13.

To define the pattern of progression of this branched trail, we analyzed the single-cell trajectory of these cells with Monocle2 [25]. In a pseudotime model, this trail started from global cluster 6, which overexpressed oligodendrocyte progenitor cell marker *OLIG1.* Then, the trail branched into 2 directions when CN-2 became CN-3, and branches 1 and 2 correspond to global clusters 10 and 12, respectively. Branch 1 exclusively expressed cilium markers (e.g., *HYDIN, FOXJ1, DNALI1,* etc.) (Fig. 5H and I), while branch 2 maintained the PN nature but showed high proliferative ability. Overexpression of *TP73* and *HYDIN* was validated by immunohistochemistry (IHC) staining experiment (Supplementary Fig. S13). Moreover, expression of the *TP73* gene at *chr1p36* increased in *chr1^amp^* cells (Fig. 5H). From previous studies, a protumorigenic activity of *TP73* has recently emerged, and especially in the context of the N-terminal truncated *TP73* isoform [26]. Furthermore, *TP73* overexpression has been reported in several tumor types, including breast cancer, melanoma, prostate cancer and neuroblastoma, and was shown to induce metastasis, chemo-resistance and other hallmarks of tumor progression that confer poor clinical outcome [27]. By RT-qPCR detection, both full length and *ΔN-TP73* existed in GS13 tumor (Supplementary Fig. S14).

In conclusion, the expression of a motile cilium signature is not an unusual event in GBM, but little is known about this particular GBM phenotype. The branched model revealed that glioma cells may undergo through different spatial destinations despite sharing similar CNV profiles.

### Characterization of the Tumor Micro Environment (TME) in Glioma

In the glioma TME, tumor associated macrophages (TAM) communicate through ligand/receptor cross-talks with tumor and non-tumor cells to promote tumor aggressiveness [28,29]. Since our single-cell platform exhibit high sensitivity with higher number of average detectable genes expressed in single cells compared to other glioma datasets (4470 genes/cell versus < 2000 genes/cell), we sought to build a ligand/receptor interaction map for the reconstruction of the most important chemoattractant relationships that exist between glioma tumor cells and TAM in the glioma TME (Fig. 6A and Supplementary Fig. S15). Overall, we detected 16 chemokine ligands and 9 receptors in 13 patients.

**Figure 6.**
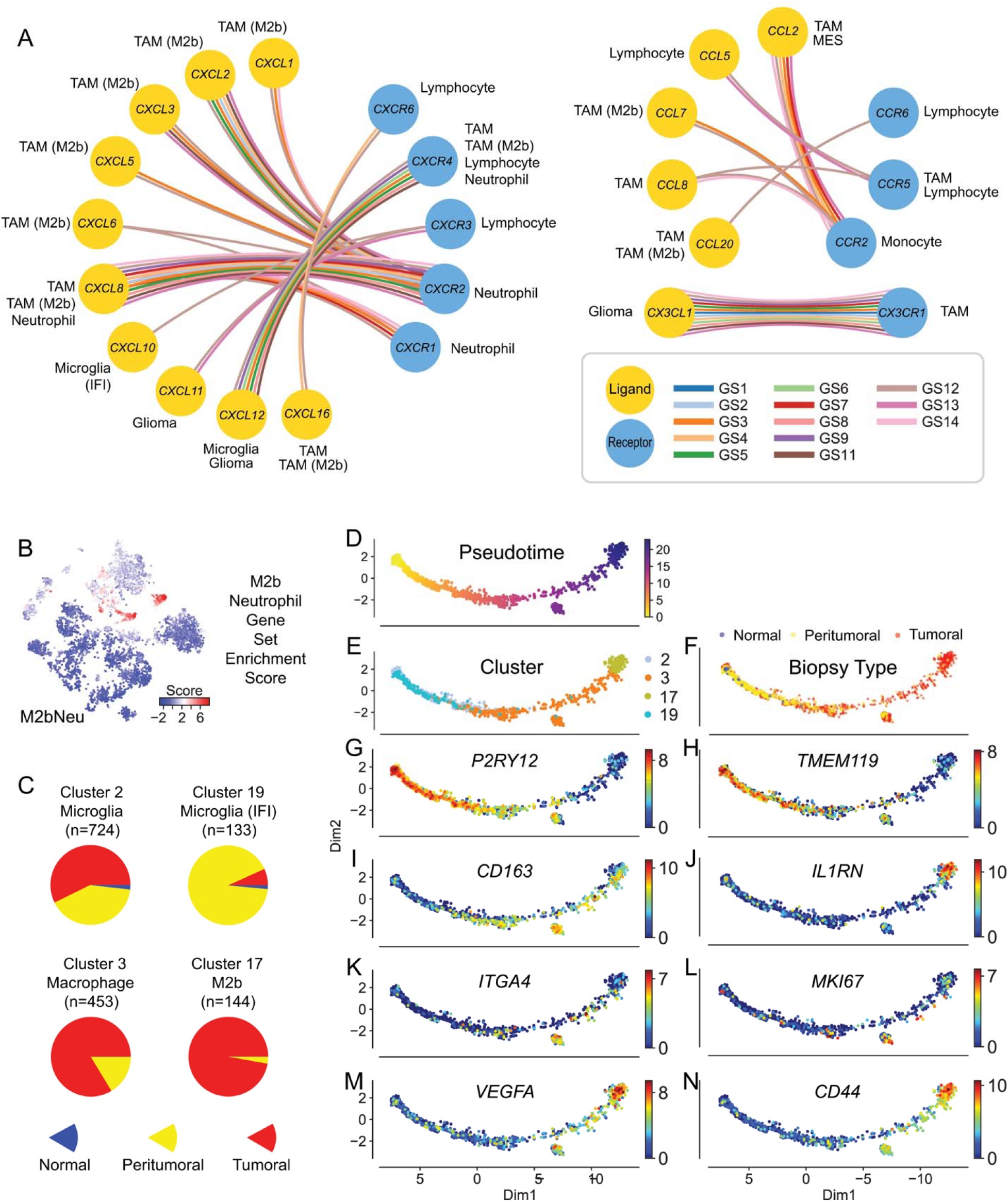
Characteristics of M2b macrophages and neutrophils and their potential in prognosis prediction using M2bNeu score. (A) Major chemokine and chemokine receptor’s relationship inside glioma tissue. (B) Dot plots showed the M2bNeu score distribution in the optimized global t-SNE map. These scores were calculated by M2bNeu genes listed in Supplementary Fig. 14C. (C) Biopsy type distribution of tumor associated microglia and macrophage cells. (D) Trajectory analysis of macrophage/microglia evolution on TAM cells from all patients, colored by (D) pseudotime, (E) optimized global t-SNE clusters, (F) biopsy types, and (G-N) marker genes expressed in the pseudotime trajectory map.

Microglia and M2a/c macrophages, which expressed the *CX3CR1* receptor, coexisted with glioma cells that expressed the *CX3CL1* ligand. Lymphocyte infiltrates expressed *CXCR3/6* and *CCR6* but the *CXCR3* ligands *CXCL9/10/11* were rarely detected in glioma samples with the exception of interferon activated microglia. These ligand-receptor pairs were previously reported to recruit tumor-infiltrating lymphocytes and inhibits tumor growth [30]. *CXCR6* ligand *CXCL16* which existed in both transmembrane and soluble form [31], was highly expressed in TAM cells, and mildly expressed in the malignant cells. Lymphocytes also expressed *CXCR4* like TAM cells, whereas the ligands were expressed in microglia cells. Their binding was reported to mediate glioma chemotaxis and regulate cell survival through activating *AKT* related pathways [32].

*CCL5/CCL8* and *CCR5* was another chemokine axis between lymphocytes and TAM. Besides inflammatory chemoattractant functions, it can also mediate NK cell activation, which promotes tumor genesis and metastasis [33]. Another receptor of *CCL8* is *CCR2*, which is expressed in monocytes, was highly expressed in the TAM cells and MES cells.

A strong impact from the *CXCL* family and related receptors was found in M2b macrophages and neutrophils (Supplementary Fig. S16A and B). *CXCL1/2/3/5/6/8* were overexpressed in M2b cells, generating a chemoattractant environment which might recruit *CXCR1^+^/CXCR2^+^* neutrophils. In glioma, M2b polarization and recruiting neutrophils have been connected with pro-tumoral functions [34–36]. Based on their common gene expression signals, we calculated an enrichment score from 38 genes (Fig. 6B and Supplementary Fig. S16C) and used this signature to deconvolute the presence of these cells from gene expression profiles of bulk tissues and predict clinical outcome. This method had been validated with TCGA glioma RNA-seq datasets (Supplementary Fig. S16D and E).

In our spatial cell distribution data, macrophages were the most abundant non-tumor cell in core biopsies, but were replaced by larger fractions of microglial cells in biopsies from the tumor margins (Fig. 6C). Thus, a switch from macrophage to microglia infiltration from the glioma core to the periphery was a general event that likely marked the microenvironmental changes. The microenvironment changes are likely to be dictated by different requirements of glioma cells as they migrate from core tumor regions to the invading front at the tumor periphery. This pattern was also recapitulated in the IVY GAP dataset [37] (Supplementary Fig. S17).

Finally, a lineage trajectory was built with TAM cells (Fig. 6D-F), showing the gradual change of three transitional states. The pattern of TAM development in glioma started first with a microglia phenotype *(P2RY12^+^/TMEM119^+^),* then it turned to polarized macrophage *(CD163^+^),* and finally converged into M2b macrophages (*IL1RN^+^*) with activated expression of strong angiogenesis signaling molecules *(VEGFA).* In the middle of this trajectory, we detected a small branch of cells expressed high levels of the bone-marrow-derived macrophage (BMDM) TAM marker *ITGA4.* These findings suggest a model whereby TAM cell polarization in glioma is the result of two independent cell sources, resident microglial cells and BMDM cells (Fig. 6G-N). As we have also been able to compare the cell fates of TAM between the different grades of glioma, we found that they exhibited lower M2 polarization in LGG samples (GS8/9/14) when compared to GBM (Supplementary Fig. S18).

## DISCUSSION

As malignant gliomas are characterized by high degree of intratumoral heterogeneity, single-cell genomic technologies have rapidly emerged as a crucial approach to disentangle glioma heterogeneity. However, most of the previous single cell RNA-seq glioma studies relied on the analysis of a single biopsy from each tumor specimen, thus lacking information on the special heterogeneity of the analyzed tumor lesions [6–9]. Although one study reported the multiregional analysis of glioma with single-cell sequencing, the number of cells analyzed in that study was very limited and could not provide a comprehensive picture of the geographical structure of glioma at the single cell level [10].

Here, we presented a comprehensive single-cell landscape of multiple subtypes of gliomas, each of which was analyzed by multi-region samplings, provided the first spatial-level analysis of the cellular states that characterize human gliomas. We designed multisector biopsies with 3D enhanced MRI model, and collected them during surgery by navigation sampling. For each biopsy, we generated and functionally annotated transcriptomes of hundreds of single tumor and non-tumor cells collected from multiple core and periphery tumor locations. Together, they provide a coherent map of the dynamic states and interactions between the different cell types that integrates the key features of glioma homeostasis at each tumor location. We found that both the number and the transcriptomic subtypes assigned to individual glioma cells frequently change dramatically between biopsies collected from different locations, even when they originated from neighboring glioma regions. We also made the unexpected observation that whereas core biopsies contained a high number of macrophages, this configuration of the core TME was replaced by a comparatively higher number of resident microglia at the glioma periphery, which represents the invading front of the tumor towards the normal brain.

As malignant glioma cells share high proliferation capacity, they readily accumulate multiple types of genetic alterations that trigger an increasing degree of aneuploidy with constant adaptation to the demands created by the growing tumor mass in relation to the TME [38]. A glioma cell CNV-driven progression trajectory we uncovered that was especially highlighted by the dynamic changes in the tumor from patient GS1 was marked by progressive loss of the astrocyte-like hallmarks of glioma cells with gain of multiple tumor cell phenotypes (loss of differentiation, competence to migrate and invade through the extracellular matrix, etc.) that together drive glioma progression and invasion of the normal brain. Another progression trajectory we uncovered was well represented in the tumor of patient GS13, in which the pseudotime trajectory produced by Monocle2 identified a drastic switch in the major tumor cell population with gain of an intriguing ciliated phenotype that likely contributes to glioma aggressiveness. These transformation models indicate that the constant rearrangement of the genome of glioma tumor cells leads to continuous gain of new capacities, all of which converge towards the acquisition of more aggressive tumor phenotypes for always more deregulated proliferation, anaplasia and invasion of the normal brain.

An important, novel finding contributed by our work is the deconvolution of the cross-talks between tumor and non-tumor cells in the glioma TME. We found that the active communications between the different cell types are primarily implemented by multiple combinations of chemokine ligands with their corresponding receptors that we have characterized within different regions of individual tumors and among the different types of glioma we studied [39]. In particular, we found that the communication between non-tumor cells was dominated by the prominent role of the *CXCL* family of chemokines and related receptors, which was especially apparent in M2b macrophages and neutrophils. We followed up on this finding and determined the enrichment score of M2b/Neutrophil cells in bulk gliomas to evaluate the consequences of the infiltration of these cell types for clinical outcome of glioma patients. The analysis, which was performed with TCGA-derived glioma RNA-seq datasets, was able to distinguish patients with divergent clinical outcome based on the predicted level of infiltration of the two cell types.

In conclusion, we used the single cell RNA-seq technology to generate an extensive complete map of the geographical molecular structure of gliomas. The trajectories of reciprocal genomic and functional changes that accompany glioma cells as they move within the tridimensional space of the tumor mass, combined with the deconvolution of the cross-talks between different cells in the glioma TME, paint an unprecedented scenario that elucidates the intratumoral heterogeneity of this lethal tumor type.

## ACKNOWLEDGEMENTS

This work was supported by National Key Research and Development Program of China (2016YFC0906000 [2016YFC0906001] to X.D.S.; 2018YFA0107601, 2017YFA0102702 to F.T.; and 2016YFC0902500 to T.J.). This work was also supported by a Grant from the biological medicine department, Beijing Municipal Science & Technology Commission (No. Z141100000214009).

## AUTHOR CONTRIBUTIONS

X.D.S., F.T., T.J. conceived the project, K.Y., Y.H., F.W. designed the experiment, K.Y. and Y.H. wrote the manuscript. F.W., Z.Q., J.C. K.W. performed multisector biopsies and tumor tissue dissection. Y.H., Q.G., W.H., X.Y.F., X.W. carried out most of the experiments. K.Y. analyzed the data. X.D.S., J.R., F.T., X.L.F., I.A. discussed and revised the manuscript. K.Y., Y.H., F.W., Q.G. contributed equally to this work.

## MATERIALS AND METHODS

### Tumor samples and multisector biopsy

Tumor samples were obtained from consenting patients after they signed an informed consent document, which was in strict observance of the ethics regulations. The use of specimens in this project was approved by the institutional review board (IRB) at Beijing Tiantan Hospital under IRB KY2018-018-02.

Peri-operative T1+gadolinium and FLAIR (fluid attenuated inversion recovery) sequences enabled the objective discrimination of the gadolinium-enhanced (CE) tumor core, nonenhanced (NE) areas, and FLAIR hyperintense tissue at the tumor margins (edema). Multiple (3–8) random MRI-localized (CE, NE, edema) biopsies measuring ~1.0 cm × 0.5 cm × 0.5 cm and at least 10 mm apart were planned before surgical debulking. During the operation, we used the Brainlab Neuronavigation interface (Brainlab) to guide the biopsy and confirm the planned radiographic localization. Specimens from each region were divided in two: one piece was immediately flash frozen in liquid nitrogen, and the other piece was used for single-cell preparation.

### Tumor tissue dissection

Immediately after sample collection, the tumors were washed with phosphate-buffered saline (PBS), fully dissociated and subjected to enzymatic dissociation with trypsin-EDTA solution. Following filtrating through a cell strainer and red blood cell lysis, the cell suspensions were used for single-cell isolation.

### Single-cell RNA-seq library construction

Single-cell RNA-seq libraries were constructed following the STRT-seq [1] protocol with minor modifications that were previously described [2,3].

Briefly, we used a mouth pipette to pick single cells into cell lysis buffer, containing barcoded reverse transcription primers (5’-TCAGACGTGTGCTCTTCCGATCTXXXXXXXXNNNNNNNNT25-3’; X is the predesigned cell-specific barcodes, and N is the unique molecular identifier (UMI)), each reverse transcribed mRNA would have an unique UMI sequence since UMI sequences were included in the reverse transcription primers. The 96 cell-specific barcodes were provided in Gene Expression Omnibus (GEO) under accession number GSE117891. The first strand cDNA was synthesized with SuperScript II reverse transcriptase in the same tube of lysed cell, and hybrid to the template switching oligo (5’-AAGCAGTGGTATCAACGCAGAGTACATrGrG+G-3’; rG is riboguanosine and +G is LNA G) immediately. Then, KAPA HiFi HotStart PCR mix and cDNA amplification primers were added to the tube to finish template switching and amplification reactions. After reverse transcription and cDNA amplification, the PCR products with 96 different cell-specific barcodes were pooled into one tube. The pooled cDNA was purified by Zymo DNA Clean & Concentrator Kit (D4013), and eluted in 50 μL H2O, then purified twice with 0.8X AMPURE XP magnetic beads to remove primers and short fragments. To tag the 3’ prime of full length cDNA with Biotin, we used 3’ biotin index primers (5’− /biotin/CAAGCAGAAGACGGCATACGAGAT/index/GTGACTGGAGTTCAGACGTGTGCTC TTCCGATC-3’) and 5’ cDNA amplification primer, amplified the cDNA for 4 cycles with KAPA HiFi HotStart PCR enzyme, then purified with 0.8X AMPURE XP beads.

To enrich the 3’ ends of cDNA, the biotin-tagged full length cDNA were sonicated with Covaris S220 machine with 300 bp program. Fragmented product were purified with Zymo DNA Clean & Concentrator Kit, and 1X AMPURE XP beads. 10 μL washed Dynabeads MyOne streptavidin C1 (Invitrogen, 65002) magnetic beads were used to enrich the biotin-tagged 3’ fragments. After binding, 100 μL 1X B&W buffer and 100 μL elution buffer were used to wash magnetic beads, then resuspend beads in 25 μL ddH_2_O. End-repair, dA-tailing and adaptor ligation were done on bead with KAPA Hyper Prep Kits (KK8505). The library amplification primers were an Illumina QP2 primer (5’-CAAGCAGAAGACGGCATACGA-3’) and a short universal primer (5’- AATGATACGGCGACCACCGAGATCTACACTCTTTCCCTACACGAC-3’). The libraries were sequenced on Illumina HiSeq 4000 platform to generate 150-bp paired-end reads.

### Whole genome DNA sequencing library construction

We extracted genomic DNA using the DNeasy Blood and Tissue Kit (Qiagen) following the manufacturer’s instructions. DNA was then fragmented to an average of 315 bp. The libraries were constructed using KAPA Hyper Prep Kits (KK8505) and sequenced on Illumina HiSeq 4000 platform.

### Single-cell RNA-seq data processing

Raw sequencing reads of single cell were obtained from pooled library data by cell-specific barcodes at the beginning of read 2, followed by trimming of the read 1 TSO sequence with an in-house script. More quality control procedures were applied to the remaining sequence of read 1; sequences containing poly-A tails, sequencing adapters, or low quality bases (N bases > 10%) were removed. Clean data were aligned to the hg38 human reference genome with hisat2 (version 2.0.5) [4]. Uniquely mapped reads on each gene were calculated with htseq-count, and PCR redundant reads were eliminated with UMI sequences in each paired read, generating a UMI count matrix of each gene in each cell [5].

In these 14 patients, we sequenced a total of 7,928 single cells, but only 6,148 cells with more than 10,000 UMIs and 2,000 detected genes were kept for downstream analysis. The median number of UMIs in the 6,148 cells was 97,294, and the median number of detected genes was 4,470.

Saturation analysis was performed to ensure sufficient data to make sure we have generated enough data to detect most of transcript in our single-cell library. For each single cell, down sampling was done by randomly extracting 10 to 90% of the original data, followed by calculating the detected gene number within this subset of sequencing data. The box plot in Supplementary Fig. S1A indicates that more than 90% of genes could be detected using only 70% of the data; thus, we had sufficient data for the subsequent analyses.

For the differential expression analysis, TPM (transcripts per million) values were calculated from the UMI matrix. Considering that most of the single cells did not have more than 1 million UMIs, we used a previously published method for normalization to avoid counting each transcript several times: we calculated the *log_2_(TPM/10+1)* instead of the traditional *log_2_(TPM+1).*

### CNV identification with whole genome DNA-seq and single-cell RNA-seq

To identify large fragment copy number variations (CNVs), we performed low-throughput whole genome sequencing with bulk cells from 42 sampling points, achieving an average sequencing depth of 0.66X. In each sample, we calculated the average depth of every 1-Mb window in all the chromosomes. To scale the average depth to the diploid level, we collected all the average window depth values and generated a histogram with 1,000 bins, in which diploid depth corresponded to the peak. Thus, all the raw average window depth data could be scaled based on a diploid baseline of 2, for a scaled average window depth of a normal diploid chromosome region equal to 2. This method could be used in most of our samples; however, as there was strong chromosome instability in patients GS3 and GS15, we had to define the ratio between the raw average window depth and the diploid average window depth based on observations of the ensemble copy number status, especially for the sex chromosomes (Supplementary Fig. S3B). Arm level CNV with copy number above 2.2 or below 1.8 were counted to identifying less CNV events region. If arm level CNV only happened in no more than 2 patients (except GS3 and GS15), variable genes that generated by Seurat in these regions were used for final PCA and t-SNE analysis. Expression heatmap of removed variable genes were showed in Supplementary Fig. S4.

For single cells, we adapted a previously described method [6,7] to estimate large fragment copy number alterations with scRNA-seq data. Since limited genes could be detected within single-cell data and we just wanted to identify CNV events at large fragment resolution, only genes with an average expression value *avg(log_2_(TPM+1))* > 1.5 were kept for analysis. As CNV events could yield a similar pattern of changes in expression level of these genes, the average expression level in a moving window containing 50 genes in both the upstream and downstream directions was calculated to estimate the local CNV value, as described in a previous paper, followed by scaling to the centered CNV matrix.

Because there was a significant difference between normal and malignant glial cells, we performed t-SNE analysis and identified a group of cells that not only composed a major proportion of cells in nontumor biopsy sites but also expressed several normal oligodendrocyte marker genes (*MOG*, *MAG*, etc.). These cells were identified as normal oligodendrocytes, and they should be free of CNVs. Therefore, we used these cells as the control group and calculated the average CNV values of the control group with a method similar to that described above, followed by the centering procedure. The final CNV value matrix of the selected genes was calculated by subtracting the centered control CNV values from the centered CNV matrix of each corresponding sliding window (Fig. 1D and Supplementary Fig. S3C).

To generate a large fragment CNV matrix for hierarchical clustering, the average CNV value of a fixed chromosome region size with the 10 Mb bin was calculated in each cell. In each patient, the CNV matrix subset containing major large-scale CNVs was used to identify clusters, and the chromosomes we used for clustering in different patients are described in Supplementary Fig. S3B.

### Cell clustering with PCA and t-SNE analysis in Seurat

The R package Seurat (version 1.4) was used to cluster the UMI count matrix of the 6,148 cells [8]. First, we selected highly variable genes with the MeanVarPlot function; genes with an average expression value between 0.25 and 4 and a dispersion value above 0.8 were selected for principal component analysis (PCA) and t-SNE analysis. Next, we performed linear dimensional reduction with PCA to convert the large number of variable genes into a set of representative variables, or principal components (PCs). The most representative PCs were retained for downstream t-SNE analysis by the jackstraw resampling test with 1,000 replicates. We used 30 PCs to find clusters and run the t-SNE analysis; the resolution parameter in the find clusters procedure was set to 1.5 by multiple trial and observation (Supplementary Fig. S5).

Within the clustered 2-dimensional t-SNE map, we observed a fragmented pattern in which cells from different patients were separated from others, despite expressing similar biomarkers (Fig. 2B). This finding was caused by the divergent CNV pattern in glioma patients. Thus, we used CNV information from bulk DNA sequencing, selected chromosome regions that contained fewer copy number alterations (Fig. 2A), and removed variable genes outside these regions. We reperformed the PCA and t-SNE analysis with this variable gene subset; 28 PC were used to identify clusters, and we generated the final global t-SNE map.

To validate the feasibility of our strategy that removes variable genes from regions with numerous CNVs, firstly, we checked the gene expression data of those removed genes, their pattern are following the optimized 25 cell clusters. We also used SCENIC to reanalyze our data [9]. SCENIC utilizes UMI counts to construct gene-regulatory networks and thus identify cell states. First, potential transcription factor (TF) targets were identified with gene co-expression information, followed by enrichment analysis of these TF motifs, thereby generating a subset of pruned modules called regulons. Then, regulons were scored by calculating the area under the recovery curve (AUC) for each single cell, and an AUC score matrix was generated. By inspecting the distribution of the AUC score of each regulon, the active status of each regulon was determined based on a cutoff value, and the original continuous AUC score matrix was transformed to a binary regulon activity matrix, representing “on” and “off” with the binary values “1” and “0”. With this matrix, we subsequently performed the t-SNE analysis and compared the results with those of our global t-SNE clusters (Fig. 2E). We used adjust rand index and adjusted mutual info score to evaluate the difference between SCENIC based clusters and our global Seurat clusters, the scores are 0.58 and 0.66, respectively. This is not a perfect score, however, after we fitted several important marker gene expression values into the SCENIC t-SNE map, we found that the SCENIC binary regulon matrix has its own characteristics. For major categories of cell types, like normal glial, TAM cell, tumor cell, SCENIC matrix could provide enough information to distinguish these cells because of their distinct transcription factor activity. However, inside these big categories, there is limited information in the binary regulon matrix to provide good performance in the clustering analysis. For example, the TAM cells and neutrophils were separated from most of the tumor cells significantly, however, interferon *(IFIT2^+^)* induced microglia mixed with normal microglia, neutrophil *(CXCR1^+^)* closely distributed with m2b macrophages *(CXCL2^+^),* and other tumor cells expressing different marker genes distributed next to each other closely, hard to distinguish their border. There is another issue during K-Means clustering, when the K-Means algorithm taking in the binary regulon matrix, activated regulons have different weight inside different cell types. For example, when we set the parameters to do the clustering under the premise that ensures the *MKI67^+^* proliferation cells or *CD44^+^* mesenchymal stromal-like cells could be properly clustered, other cell types would be over-clustered or underclustered. We took several marker gene expression data into consideration, merged both SCENIC clusters and Seurat clusters into 11 major cell types based on the gene expression status described in the in Supplementary Fig. S7F and G. Now, the adjust rand index and adjusted mutual info scores are 0.76 and 0.76, which could support the conclusion that those two dimensional reduction methods have a similar outcome.

### Identification of differentially expressed genes and functional enrichment analysis

To understand the cell type and functional characteristics of each cell cluster, the Seurat function FindAllMarkers was used to perform a differential expression analysis with a likelihood-ratio test and argument: only.pos = TRUE, min.pct = 0.25, thresh.use = 0.25. AUC value were also calculated with roc test. For each cluster, AUC value of each gene were sorted in descending order. Genes with a lower adjusted p-value or higher AUC value were highly expressed in this cluster and represent the function and morphological characteristics of the corresponding cluster.

In the patient sub-CNV analysis, differentially expressed (DE) genes were identified among 2 or 3 groups of cells. For these cells, we use a t-test between each of 2 groups of cells; genes that met the condition of log_2_FC > 1 and *P*-value < 0.01 were kept as significant DE genes.

Marker genes of clusters (AUC>0.7) and significant DE genes between specified cell groups were used to deduce function. We performed functional enrichment analysis with Metascape (http://metascape.org) and the R package ‘clusterProfiler’ [10].

### Single-cell trajectory analysis

During tumor development, malignant cells transition from one state to another, with differential expression of various genes. Thus, RNA-seq data for a group of related single cells could be used to deduce the relationships among these cells and construct a robust track of their changes in pseudotime. The R package Monocle2 was applied to estimate the developmental pseudotime of all malignant cells in an individual patient [11]. For each patient, we followed the standard monocle document and identified DE genes that defined progress towards the UMI count matrix of malignant cells. Next, the function reduceDimension with the DDRTree method was applied to reduce the dimensions, and the cells were then ordered along the trajectory using the orderCells function. Since Monocle produces a linear trajectory, and the direction of the pseudotime trajectory was unknown, we manually set the pseudotime direction based on other information such as CNV changes and the expression levels of progenitor or malignant marker genes.

### Gene set enrichment score calculation

We adopted the method described in published work [12] to calculate the enrichment score of a given gene set. For gene *j* in cell *i,* expression levels were first quantified as *E_i,j_* = log_2_[(*TPM_i,j_* / 10) + 1] and then centered on the relative expression (*Er_i,j_*): *Er_i,j_* = *E_i,j_* – average[*E*_*i*,1…*n*_].

To calculate the enrichment score of a given gene set, we introduced a control gene set derived from the given set. Expression levels were sorted and divided into 25 bins; then, 100 genes were randomly selected for each gene from the same expression bin. For a set of genes (*G_j_*) the score of this gene set in cell i was calculated based on the expression level of a control gene set: *SC_j_*(*i*) = average[*Er*(*G_j,i_*)] – average[*Er*(*G_jcont,i_*)].

In our analysis, we collected gene lists related to *“HALLMARK EPITHELIAL MESENCHYMAL TRANSITINSITION’* and *“HARRIS BRAIN CANCER PROGENITORS”* from the Molecular Signatures Database [13–15].

### M2b macrophage/neutrophil marker identification and survival analysis with TCGA glioma datasets

M2b macrophages and neutrophils are important factors that reflect the status and degree of inflammation and malignancy; thus, estimating the cellular constituents using bulk tumor expression data could be valuable for clinical curation. In our PCA analysis of nonmalignant cells, we found that PC2 genes perfectly screened out M2b macrophages and neutrophils. Therefore, we selected the top 200 PC2 genes, and hierarchical clustering helped us identify 78 marker genes that were highly expressed in M2b macrophages and neutrophils.

The level 3 TCGA glioblastoma and low grade glioma RNAseqV2 data were downloaded from the GDC data portal of the National Cancer Institute website. After a data cleaning procedure, TPM matrix and clinical information for 152 GBM and 502 LGG samples were prepared. Although the 78 marker genes have a similar expression pattern in our cell type expression profile, consistency among large populations would be an important element affecting their representativeness of M2b/neutrophil components in bulk tumor tissue. Therefore, we removed marker genes not generally correlated by calculating a Pearson correlation matrix of these 78 genes with both the GBM and LGG matrices, and genes with an average correlation value above 0.2 were retained. Thirty-eight genes were used to calculate the *M2bNeu* score (Supplementary Fig. S16C).

### Isolation and culturation of primary glioma cells

Tumor samples were washed and dissociated enzymatically as previously described [16]. Then, single cell suspension was cultured in neurobasal medium containing 20 ng/mL basic fibroblast growth factor *(bFGF),* 20 ng/mL of epidermal growth factor (*EGF*), 2% B27 and 10 ug/mL heparin. The tumor spheres were collected and digested with accutase for passage.

### Immunohistochemical analysis

Immunohistochemical analysis were conducted using paraffin section specimens from patients. Sections were incubated with antibodies *(TP73:* Abcam ab40658, *HYDIN:* Abcam ab221643) overnight at 4°C, and staining was performed using diaminobenzidine staining.

## DATA AVAILABILITY

Both the single-cell RNA-seq data and bulk DNA-seq data we used in this study were submitted to Sequence Read Archive (SRA) under accession number SRP153120. Barcode sequences and prepared gene expression count matrix were also submitted to the Gene Expression Omnibus (GEO) under accession number GSE117891. Other information related to this study is available from the corresponding authors upon reasonable request.

## REFERENCES

1. Vogelstein B, Papadopoulos N, Velculescu VE, Zhou S, Diaz LA, Kinzler KW. Cancer Genome Landscapes. Science (80-) 2013; 339:1546–1558.

2. Snuderl M, Fazlollahi L, Le LP, et al. Mosaic amplification of multiple receptor tyrosine kinase genes in glioblastoma. Cancer Cell 2011; 20:810–7.

3. Meyer M, Reimand J, Lan X, et al. Single cell-derived clonal analysis of human glioblastoma links functional and genomic heterogeneity. Proc Natl Acad Sci 2015; 112:851–856.

4. Sottoriva A, Spiteri I, Piccirillo SGM, et al. Intratumor heterogeneity in human glioblastoma reflects cancer evolutionary dynamics. Proc Natl Acad Sci USA 2013; 110:4009–14.

5. Suzuki H, Aoki K, Chiba K, et al. Mutational landscape and clonal architecture in grade II and III gliomas. Nat Genet 2015; 47:458–468.

6. Patel AP, Tirosh I, Trombetta JJ, et al. Single-cell RNA-seq highlights intratumoral heterogeneity in primary glioblastoma. Science 2014; 344:1396–401.

7. Tirosh I, Venteicher AS, Hebert C, et al. Single-cell RNA-seq supports a developmental hierarchy in human oligodendroglioma. Nature 2016; 539:309–313.

8. Venteicher AS, Tirosh I, Hebert C, et al. Decoupling genetics, lineages, and microenvironment in IDH-mutant gliomas by single-cell RNA-seq. Science 2017; 355.

9. Darmanis S, Sloan SA, Croote D, et al. Single-Cell RNA-Seq Analysis of Infiltrating Neoplastic Cells at the Migrating Front of Human Glioblastoma. Cell Rep 2017; 21:1399–1410.

10. Lee J-K, Wang J, Sa JK, et al. Spatiotemporal genomic architecture informs precision oncology in glioblastoma. Nat Genet 2017; 49:594–599.

11. Neftel C, Laffy J, Filbin MG, et al. An Integrative Model of Cellular States, Plasticity, and Genetics for Glioblastoma. Cell 2019; 178:835–849.e21.

12. Dong J, Hu Y, Fan X, et al. Single-cell RNA-seq analysis unveils a prevalent epithelial/mesenchymal hybrid state during mouse organogenesis. Genome Biol 2018; 19:31.

13. Gao S, Yan L, Wang R, et al. Tracing the temporal-spatial transcriptome landscapes of the human fetal digestive tract using single-cell RNA-sequencing. Nat Cell Biol 2018; 20:721–734.

14. Aibar S, González-Blas CB, Moerman T, et al. SCENIC: Single-cell regulatory network inference and clustering. Nat Methods 2017; 14:1083–1086.

15. Peschl P, Bradl M, Höftberger R, Berger T, Reindl M. Myelin Oligodendrocyte Glycoprotein: Deciphering a Target in Inflammatory Demyelinating Diseases. Front Immunol 2017; 8:529.

16. Zhan C, Yan L, Wang L, et al. Identification of immunohistochemical markers for distinguishing lung adenocarcinoma from squamous cell carcinoma. J Thorac Dis 2015; 7:1398–405.

17. Mei F, Wang H, Liu S, et al. Stage-Specific Deletion of Olig2 Conveys Opposing Functions on Differentiation and Maturation of Oligodendrocytes. J Neurosci 2013; 33:8454–8462.

18. Saadoun S, Papadopoulos MC. Aquaporin-4 in brain and spinal cord oedema. Neuroscience 2010; 168:1036–46.

19. Verhaak RGW, Hoadley KA, Purdom E, et al. Integrated Genomic Analysis Identifies Clinically Relevant Subtypes of Glioblastoma Characterized by Abnormalities in PDGFRA, IDH1, EGFR, and NF1. Cancer Cell 2010; 17:98–110.

20. Satir P, Christensen ST. Structure and function of mammalian cilia. Histochem Cell Biol 2008; 129:687–693.

21. Yu X, Ng CP, Habacher H, Roy S. Foxj1 transcription factors are master regulators of the motile ciliogenic program. Nat Genet 2008; 40:1445–53.

22. Casasent AK, Schalck A, Gao R, et al. Multiclonal Invasion in Breast Tumors Identified by Topographic Single Cell Sequencing. Cell 2018; 172:205–217.e12.

23. Wang L, Fan J, Francis JM, et al. Integrated single-cell genetic and transcriptional analysis suggests novel drivers of chronic lymphocytic leukemia. Genome Res 2017; 27:1300–1311.

24. Ling S, Hu Z, Yang Z, et al. Extremely high genetic diversity in a single tumor points to prevalence of non-Darwinian cell evolution. Proc Natl Acad Sci U S A 2015; 112:E6496–505.

25. Trapnell C, Cacchiarelli D, Grimsby J, et al. The dynamics and regulators of cell fate decisions are revealed by pseudotemporal ordering of single cells. Nat Biotechnol 2014; 32:381–386.

26. Stiewe T, Tuve S, Peter M, Tannapfel A, Elmaagacli AH, Pützer BM. Quantitative TP73 transcript analysis in hepatocellular carcinomas. Clin Cancer Res 2004; 10:626–33.

27. Buhlmann S, Pützer BM. DNp73 a matter of cancer: mechanisms and clinical implications. Biochim Biophys Acta 2008; 1785:207–16.

28. Quail DF, Joyce JA. The Microenvironmental Landscape of Brain Tumors. Cancer Cell 2017; 31:326–341.

29. Cohen M, Giladi A, Gorki A-D, et al. Lung Single-Cell Signaling Interaction Map Reveals Basophil Role in Macrophage Imprinting. Cell 2018; 175:1031–1044.e18.

30. Groom JR, Luster AD. CXCR3 in T cell function. Exp Cell Res 2011; 317:620–31.

31. Deng L, Chen N, Li Y, Zheng H, Lei Q. CXCR6/CXCL16 functions as a regulator in metastasis and progression of cancer. Biochim Biophys Acta 2010; 1806:42–9.

32. Zhou Y, Larsen PH, Hao C, Yong VW. CXCR4 is a major chemokine receptor on glioma cells and mediates their survival. J Biol Chem 2002; 277:49481–7.

33. Arango Duque G, Descoteaux A. Macrophage cytokines: involvement in immunity and infectious diseases. Front Immunol 2014; 5:491.

34. Balkwill FR. The chemokine system and cancer. J Pathol 2012; 226:148–157.

35. Qian B-Z, Pollard JW. Macrophage diversity enhances tumor progression and metastasis. Cell 2010; 141:39–51.

36. Turner MD, Nedjai B, Hurst T, Pennington DJ. Cytokines and chemokines: At the crossroads of cell signalling and inflammatory disease. Biochim Biophys Acta - Mol Cell Res 2014; 1843:2563–2582.

37. Puchalski RB, Shah N, Miller J, et al. An anatomic transcriptional atlas of human glioblastoma. Science 2018; 360:660–663.

38. Wu C-I, Wang H-Y, Ling S, Lu X. The Ecology and Evolution of Cancer: The Ultra-Microevolutionary Process. Annu Rev Genet 2016; 50:347–369.

39. Mollica Poeta V, Massara M, Capucetti A, Bonecchi R. Chemokines and Chemokine Receptors: New Targets for Cancer Immunotherapy. Front Immunol 2019; 10:379.

## REFERENCES

1. Islam S, Kjällquist U, Moliner A, et al. Characterization of the single-cell transcriptional landscape by highly multiplex RNA-seq. Genome Res 2011; 21:1160–7.

2. Dong J, Hu Y, Fan X, et al. Single-cell RNA-seq analysis unveils a prevalent epithelial/mesenchymal hybrid state during mouse organogenesis. Genome Biol 2018; 19:31.

3. Gao S, Yan L, Wang R, et al. Tracing the temporal-spatial transcriptome landscapes of the human fetal digestive tract using single-cell RNA-sequencing. Nat Cell Biol 2018; 20:721–734.

4. Kim D, Langmead B, Salzberg SL. HISAT: a fast spliced aligner with low memory requirements. Nat Methods 2015; 12:357–60.

5. Anders S, Pyl PT, Huber W. HTSeq--a Python framework to work with high-throughput sequencing data. Bioinformatics 2015; 31:166–9.

8. Satija R, Farrell JA, Gennert D, Schier AF, Regev A. Spatial reconstruction of single-cell gene expression data. Nat Biotechnol 2015; 33:495–502.

9. Aibar S, González-Blas CB, Moerman T, et al. SCENIC: Single-cell regulatory network inference and clustering. Nat Methods 2017; 14:1083–1086.

10. Yu G, Wang L-G, Han Y, He Q-Y. clusterProfiler: an R package for comparing biological themes among gene clusters. OMICS 2012; 16:284–7.

11. Trapnell C, Cacchiarelli D, Grimsby J, et al. The dynamics and regulators of cell fate decisions are revealed by pseudotemporal ordering of single cells. Nat Biotechnol 2014; 32:381–386.

12. Venteicher AS, Tirosh I, Hebert C, et al. Decoupling genetics, lineages, and microenvironment in IDH-mutant gliomas by single-cell RNA-seq. Science 2017; 355.

13. Subramanian A, Tamayo P, Mootha VK, et al. Gene set enrichment analysis: a knowledge-based approach for interpreting genome-wide expression profiles. Proc Natl Acad Sci U S A 2005; 102:15545–50.

14. Liberzon A, Birger C, Thorvaldsdóttir H, Ghandi M, Mesirov JP, Tamayo P. The Molecular Signatures Database (MSigDB) hallmark gene set collection. Cell Syst 2015; 1:417–425.

15. Harris MA, Yang H, Low BE, et al. Cancer stem cells are enriched in the side population cells in a mouse model of glioma. Cancer Res 2008; 68:10051–10059.

16. Wu F, Hu P, Li D, et al. RhoGDIα suppresses self-renewal and tumorigenesis of glioma stem cells. Oncotarget 2016; 7:61619–61629.

