## Supplementary for "Surveying Brain Tumor Heterogeneity by Single-Cell RNA Sequencing of Multi-sector Biopsies"

### Supplementary Information

#### Supplementary Figures

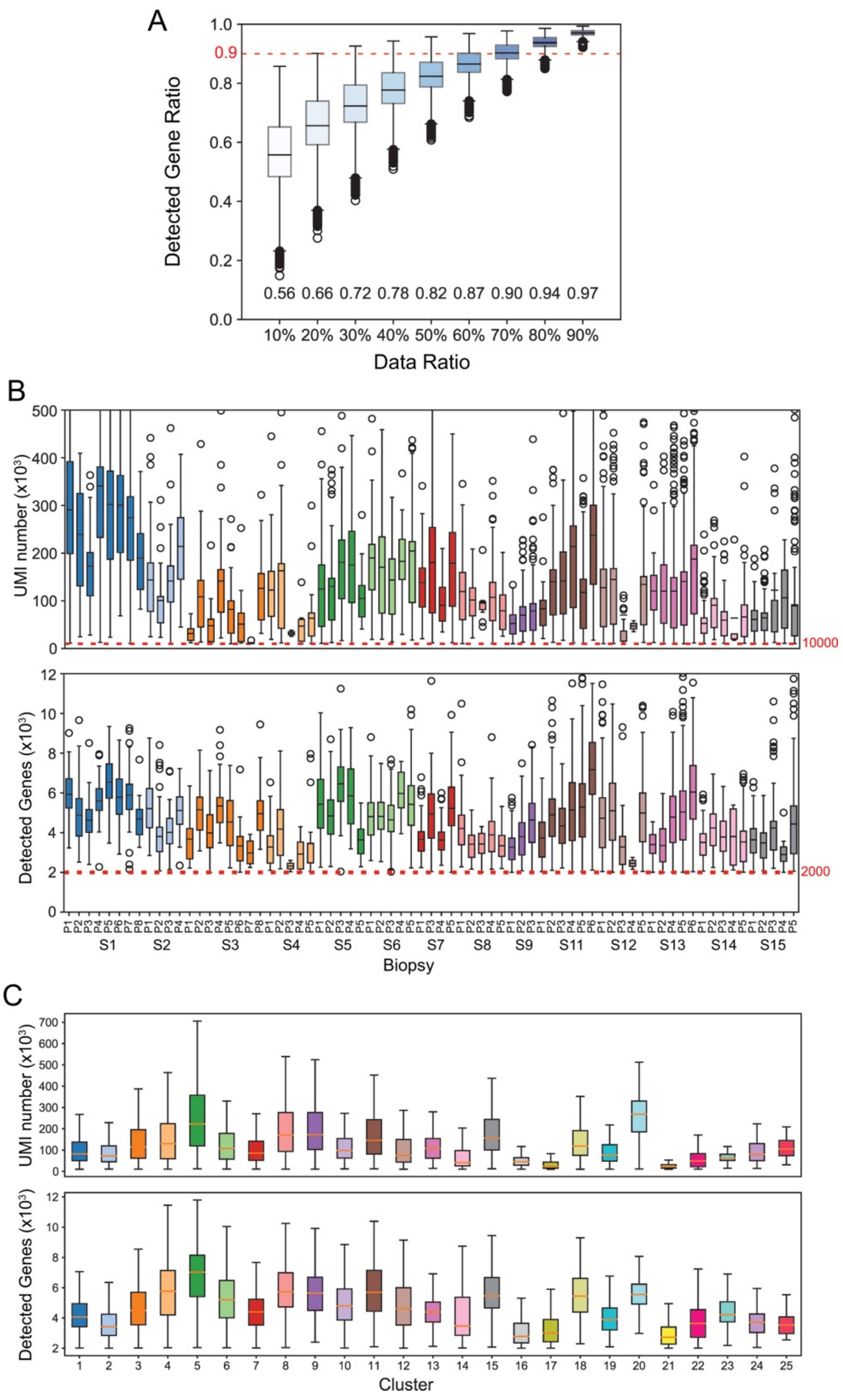

**Supplementary Fig. S1.** General Information of Experiment Procedure and Generated Data.

(A) Saturation analysis. Down sampling were done by reduce raw sequencing data to 10, 20, 30, 40, 50, 60, 70, 80 and 90% of its original data, thus calculated a series of ratio that describe detected genes proportion compared to using all raw data. (B) Boxplot of data summary, grouped by tumor biopsies, described UMI number and detected gene number of all quality filtered cells. (C) Boxplot of data summary, grouped by optimized global t-SNE clusters, described UMI number and detected gene number of all quality filtered cells.

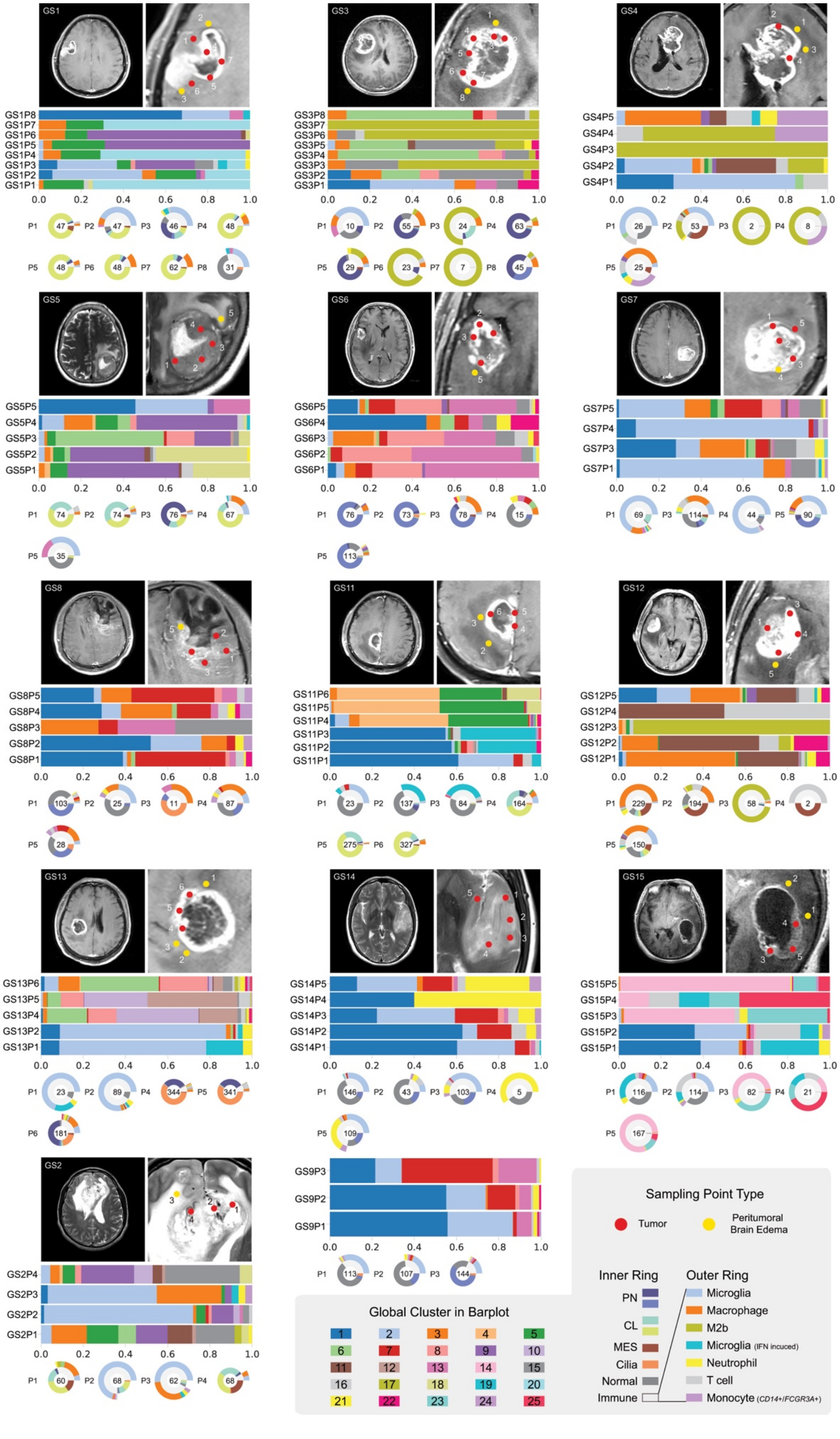
GS1
GS3
GS4
GS5
GS6
GS7
GS8
GS11
GS12
GS13
GS14
GS15
GS2
GS9P3
GS9P2
GS9P1
0.0 0.2 0.4 0.6 0.8 1.0
Sampling Point Type
Tumor
Peritumoral Brain Edema
Inner Ring
PN
CL
MES
Cilia
Normal
Immune
Outer Ring
Microglia
Macrophage
M2b
Microglia (IFN incuced)
Neutrophil
T cell
Monocyte (CD14+/FCGR3A+)
Global Cluster in Barplot
1 2 3 4 5
6 7 8 9 10
11 12 13 14 15
16 17 18 19 20
21 22 23 24 25

**Supplementary Fig. S2.** Medical Imaging and Cell Components of All Sampling Points in Each Patient.

Red and yellow dots marked locations of tumoral and peritumoral sampling point in the MRI image. Most of these MRI images were gadolinium enhanced except GS2. Bar plots illustrated cell components of each sampling points, representing 25 clusters in global t-SNE map, in accordance with the color code in (Fig. 1A and C). Ring plot illustrated cell components referred to TCGA 4 subtype classification (inner ring) and immune related cell types (outer ring).

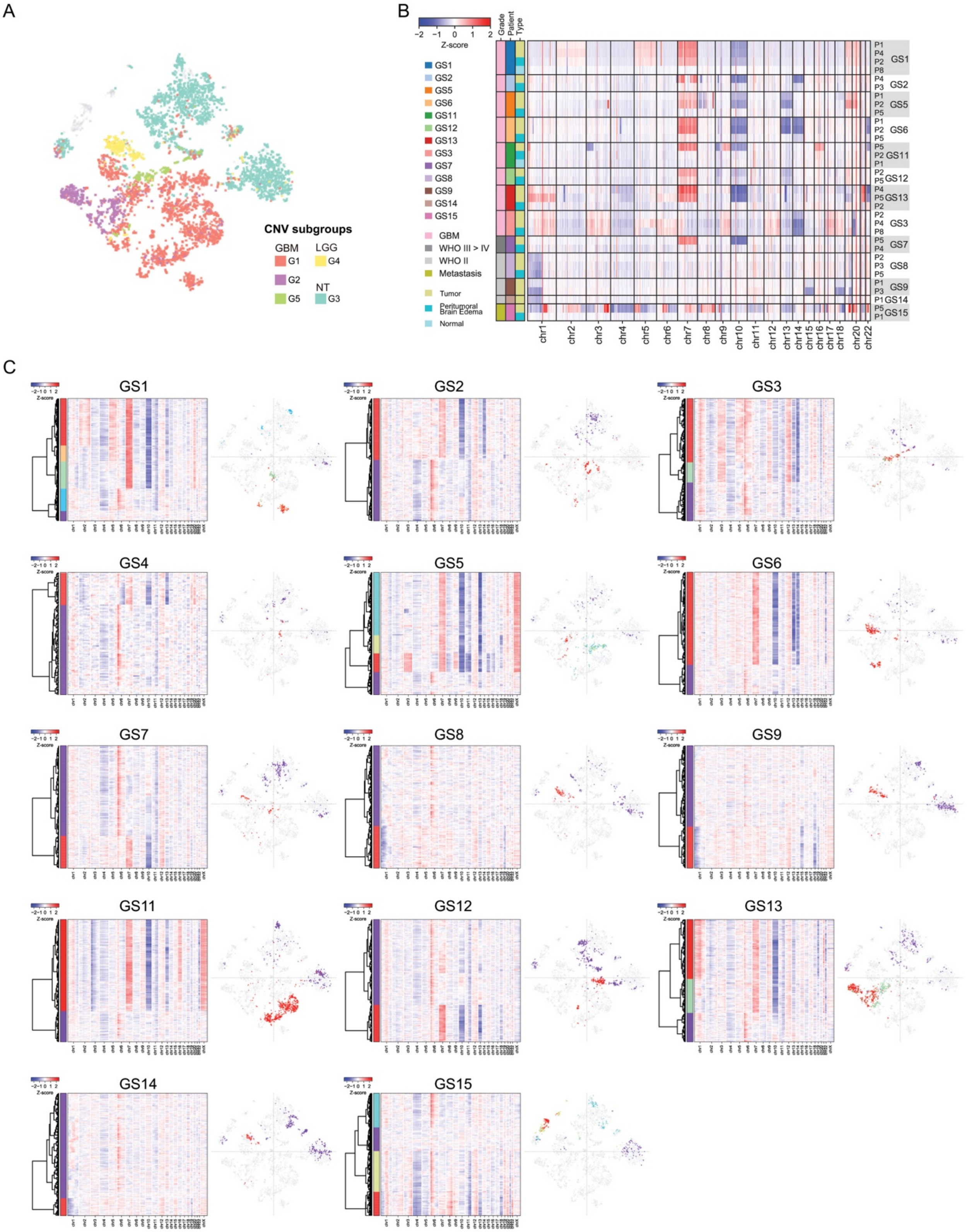
A
B
C
CNV subgroups
GBM
G1
G2
G5
LGG
G4
NT
G3
Z-score
Grade
Patient
Type
GS1
GS2
GS5
GS6
GS11
GS12
GS13
GS3
GS7
GS8
GS9
GS14
GS15
GBM
WHO III > IV
WHO II
Metastasis
Tumor
Peritumoral Brain Edema
Normal
chr1
chr2
chr3
chr4
chr5
chr6
chr7
chr8
chr9
chr10
chr11
chr12
chr13
chr14
chr15
chr16
chr17
chr18
chr20
chr22
GS1
GS2
GS3
GS4
GS5
GS6
GS7
GS8
GS9
GS11
GS12
GS13
GS14
GS15

**Supplementary Fig. S3.** CNV Information at Bulk and Single-Cell Level.
(A) CNV subtype distributions in t-SNE map. (B) Bulk CNV status of sampling points by low depth whole genome sequencing. (C) Heatmaps demonstrated RNA-seq derived CNV pattern at single-cell level, each row is a single-cell, and hierarchical clustering divided these cells into several groups. Corresponding distributions were mapped in the global t-SNE coordinates.

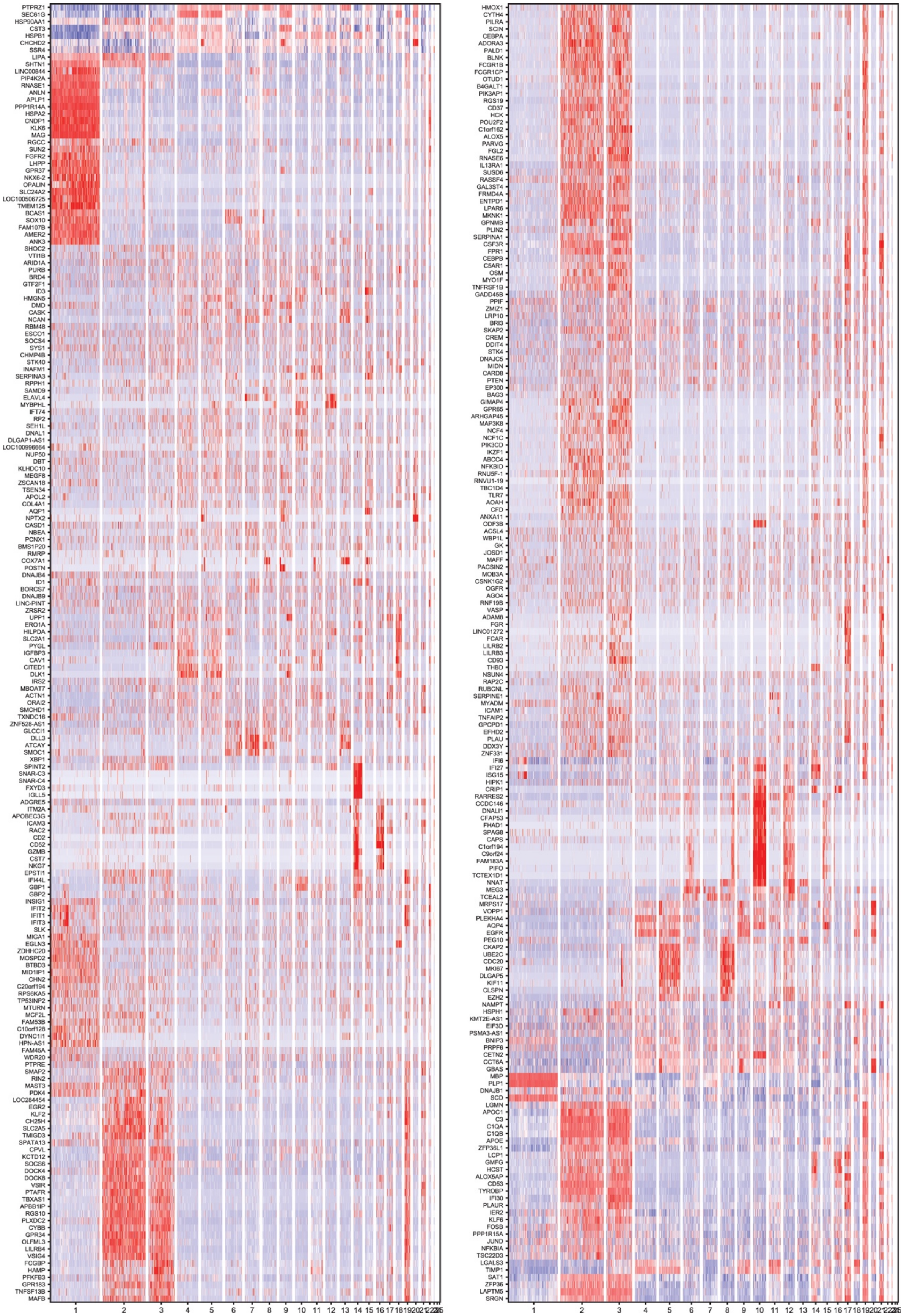
PTPRZ1
SEC61G
HSP90AA1
CST3
HSPB1
CHCHD2
SSR4
LIPA
SHTN1
LINC00844
PIP4K2A
RNASE1
ANLN
APLP1
PPP1R14A
HSPA2
CNDP1
KLK6
MAG
RGCC
SUN2
FGFR2
LHPP
GPR37
NKX6-2
OPALIN
SLC24A2
LOC100506725
TMEM125
BCAS1
SOX10
FAM107B
AMER2
ANK3
SHOC2
VTI1B
ARID1A
PURB
BRD4
GTF2F1
ID3
HMGN5
DMD
CASK
NCAN
RBM48
ESCO1
SOCS4
SYS1
CHMP4B
STK40
INAFM1
SERPINA3
RPPH1
SAMD9
ELAVL4
MYBPHL
IFT74
RP2
SEH1L
DNAL1
DLGAP1-AS1
LOC100996664
NUP50
DBT
KLHDC10
MEGF8
ZSCAN18
TSEN34
APOL2
COL4A1
AQP1
NPTX2
CASD1
NBEA
PCNX1
BMS1P20
RMRP
COX7A1
POSTN
DNAJB4
ID1
BORCS7
DNAJB9
LINC-PINT
ZRSR2
UPP1
ERO1A
HILPDA
SLC2A1
PYGL
IGFBP3
CAV1
CITED1
DLK1
IRS2
MBOAT7
ACTN1
ORAI2
SMCHD1
TXNDC16
ZNF528-AS1
GLCCI1
DLL3
ATCAY
SMOC1
XBP1
SPINT2
SNAR-C3
SNAR-C4
FXYD3
IGLL5
ADGRE5
ITM2A
APOBEC3G
ICAM3
RAC2
CD2
CD52
GZMB
CST7
NKG7
EPSTI1
IFI44L
GBP1
GBP2
INSIG1
IFIT2
IFIT1
IFIT3
SLK
MIGA1
EGLN3
ZDHHC20
MOSPD2
BTBD3
MID1IP1
CHN2
C20orf194
RPS6KA5
TP53INP2
MTURN
MCF2L
FAM53B
C10orf128
DYNC1I1
HPN-AS1
FAM45A
WDR20
PTPRE
SMAP2
RIN2
MAST3
PDK4
LOC284454
EGR2
KLF2
CH25H
SLC2A5
TMIGD3
SPATA13
CPVL
KCTD12
SOCS6
DOCK4
DOCK8
VSIR
PTAFR
TBXAS1
APBB1IP
RGS10
PLXDC2
CYBB
GPR34
OLFML3
LILRB4
VSIG4
FCGBP
HAMP
PFKFB3
GPR183
TNFSF13B
MAFB
1 2 3 4 5 6 7 8 9 10 11 12 13 14 15 16 17 18 19 20 21
HMOX1
CYTH4
PILRA
SCIN
CEBPA
ADORA3
PALD1
BLNK
FCGR1B
FCGR1CP
OTUD1
B4GALT1
PIK3AP1
RGS19
CD37
HCK
POU2F2
C1orf162
ALOX5
PARVG
FGL2
RNASE6
IL13RA1
SUSD6
RASSF4
GAL3ST4
FRMD4A
ENTPD1
LPAR6
MKNK1
GPNMB
PLIN2
SERPINA1
CSF3R
FPR1
CEBPB
C5AR1
OSM
MYO1F
TNFRSF1B
GADD45B
PPIF
ZMIZ1
LRP10
BRI3
SKAP2
CREM
DDIT4
STK4
DNAJC5
MIDN
CARD8
PTEN
EP300
BAG3
GIMAP4
GPR65
ARHGAP45
MAP3K8
NCF4
NCF1C
PIK3CD
IKZF1
ABCC4
NFKBID
RNU5F-1
RNVU1-19
TBC1D4
TLR7
AOAH
CFD
ANXA11
ODF3B
ACSL4
WBP1L
GK
JOSD1
MAFF
PACSIN2
MOB3A
CSNK1G2
OGFR
AGO4
RNF19B
VASP
ADAM8
FGR
LINC01272
FCAR
LILRB2
LILRB3
CD93
THBD
NSUN4
RAP2C
RUBCNL
SERPINE1
MYADM
ICAM1
TNFAIP2
GPCPD1
EFHD2
PLAU
DDX3Y
ZNF331
IFI6
IFI27
ISG15
HIPK1
CRIP1
RARRES2
CCDC146
DNALI1
CFAP53
FHAD1
SPAG8
CAPS
C1orf194
C9orf24
FAM183A
PIFO
TCTEX1D1
NNAT
MEG3
TCEAL2
MRPS17
VOPP1
PLEKHA4
AQP4
EGFR
PEG10
CKAP2
UBE2C
CDC20
MKI67
DLGAP5
KIF11
CLSPN
EZH2
NAMPT
HSPH1
KMT2E-AS1
EIF3D
PSMA3-AS1
BNIP3
PRPF6
CETN2
CCT6A
GBAS
MBP
PLP1
DNAJB1
SCD
LGMN
APOC1
C3
C1QA
C1QB
APOE
ZFP36L1
LCP1
GMFG
HCST
ALOX5AP
CD53
TYROBP
IFI30
PLAUR
IER2
KLF6
FOSB
PPP1R15A
JUND
NFKBIA
TSC22D3
LGALS3
TIMP1
SAT1
ZFP36
LAPTM5
SRGN
1 2 3 4 5 6 7 8 9 10 11 12 13 14 15 16 17 18 19 20 21

**Supplementary Fig. S4.** Gene expression heatmap of genes removed in the PCA and t-SNE analysis. These genes are located on chromosome regions that containing major CNV events.

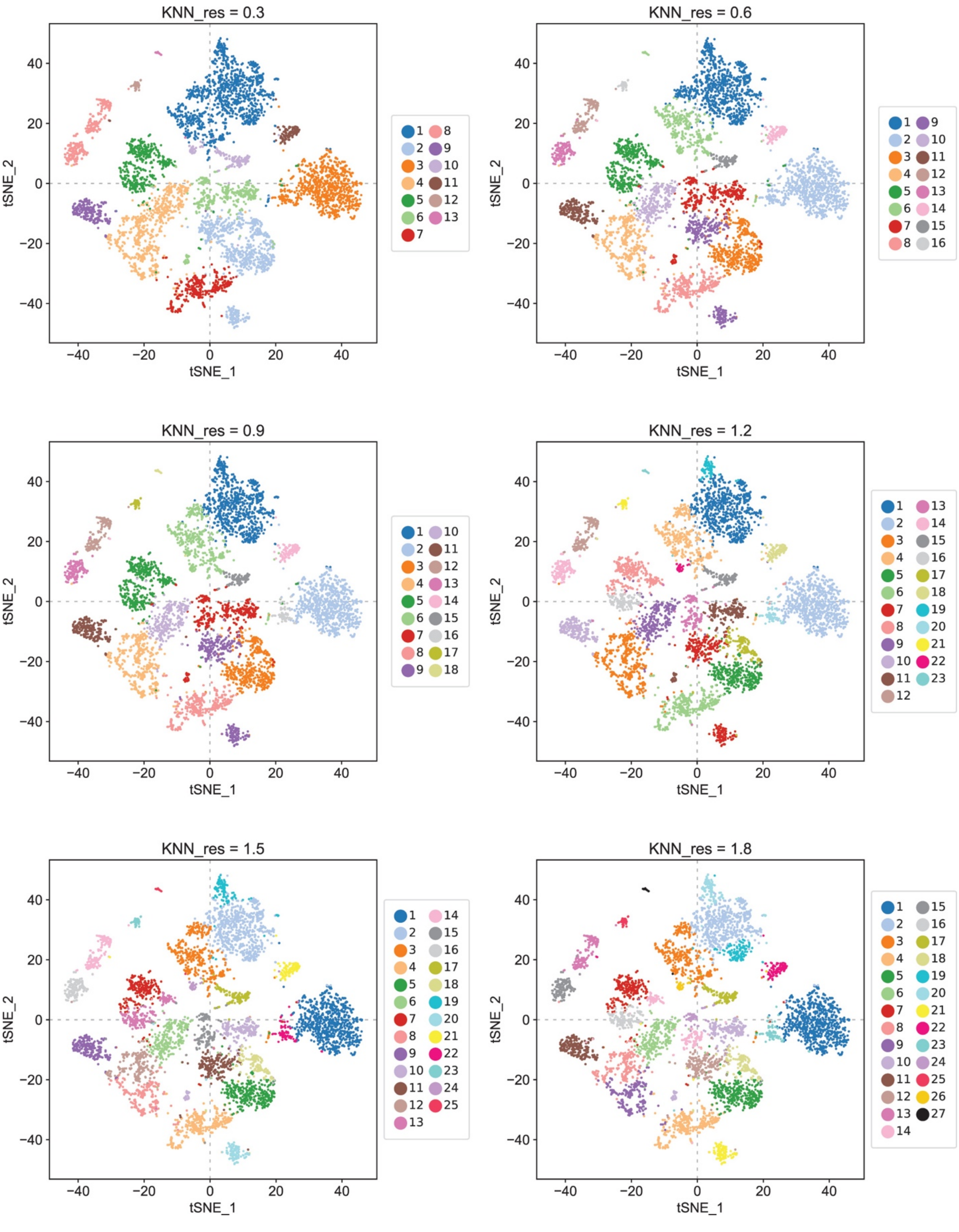

KNN_res = 0.3
KNN_res = 0.6
KNN_res = 0.9
KNN_res = 1.2
KNN_res = 1.5
KNN_res = 1.8
tSNE_1
tSNE_2

**Supplementary Fig. S5.** KNN clusters obtained with variable values of the resolution parameter.

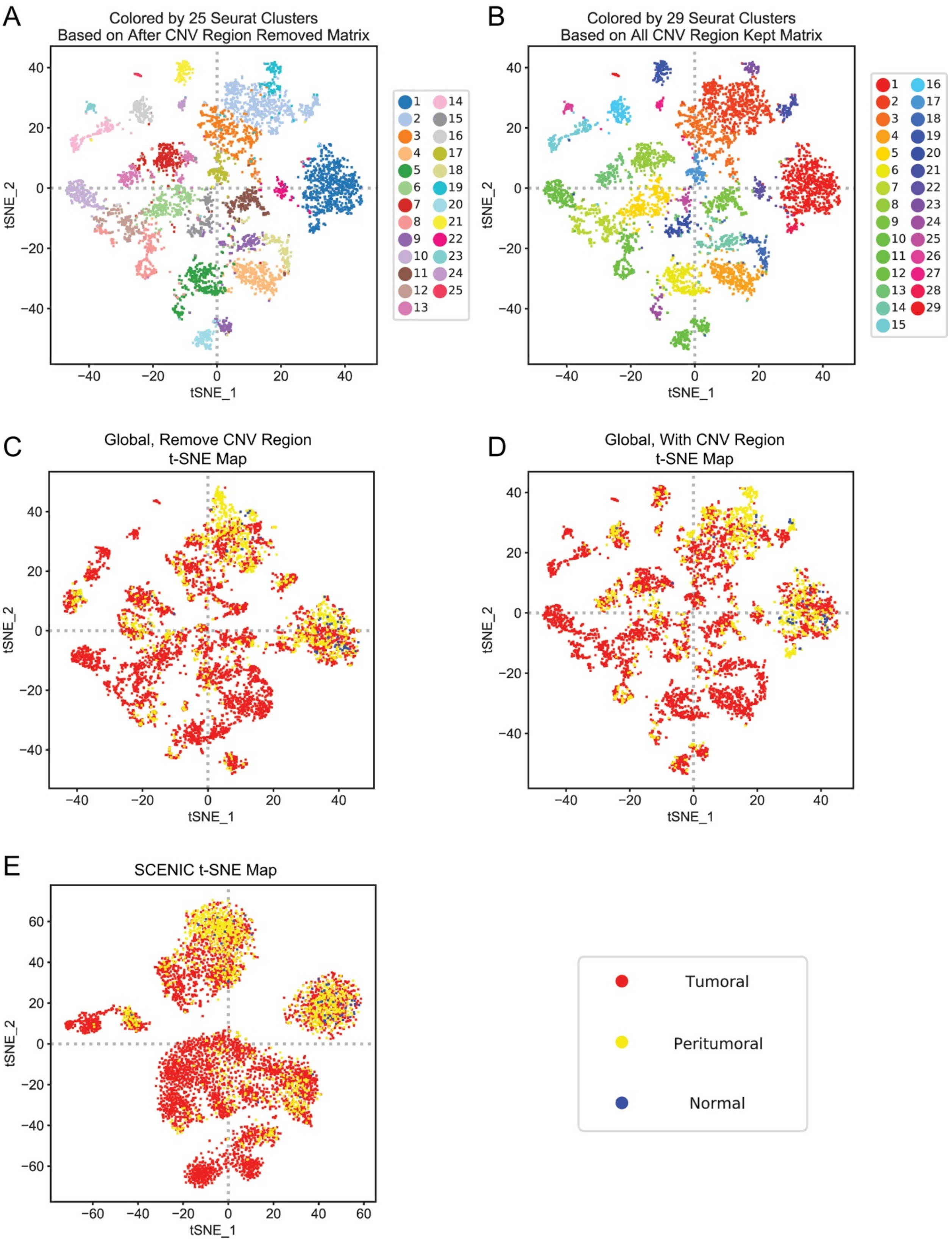

A
Colored by 25 Seurat Clusters
Based on After CNV Region Removed Matrix
B
Colored by 29 Seurat Clusters
Based on All CNV Region Kept Matrix
C
Global, Remove CNV Region
t-SNE Map
D
Global, With CNV Region
t-SNE Map
E
SCENIC t-SNE Map
tSNE_1
tSNE_2
Tumoral
Peritumoral
Normal

**Supplementary Fig. S6.** Comprehensive single cell information in different dimension reduction coordinates.

(A-B), Coordinates based on all variable genes Seurat t-SNE analysis, colored by (A) optimized global t-SNE clusters and (B) all variable genes derived original global t-SNE clusters. (C-F) Cells colored by biopsy types, mapped on (C) optimized Seurat t-SNE coordinates, (D) all variable genes derived original Seurat t-SNE coordinates, (E) SCENIC regulon based t-SNE.

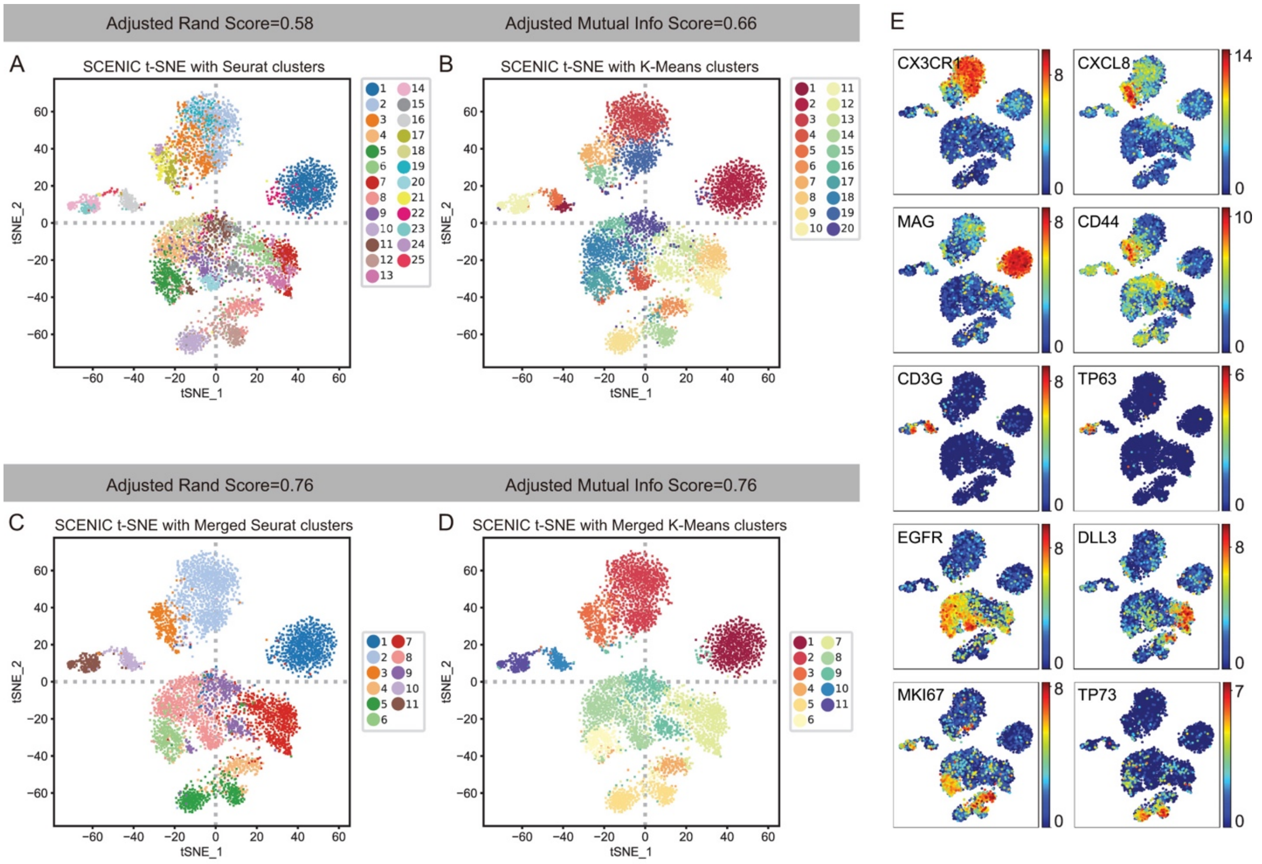


F

| Merged Seurat Clusters | Original Seurat Clusters | Marker |
|---|---|---|
| 1 | 1, 22 | MAG⁺ |
| 2 | 2, 3, 19 | CX3CR1⁺ |
| 3 | 17, 21, 24 | CXCL8⁺ |
| 4 | 8 | DLL3⁺, MKI67⁺ |
| 5 | 10, 12 | TP73⁺ |
| 6 | 5 | EGFR⁺, MKI67⁺ |
| 7 | 6, 7, 13 | DLL3⁺, MKI67⁻ |
| 8 | 4, 9, 18, 20 | EGFR⁺, MKI67⁻ |
| 9 | 11, 15 | CD44⁺, CXCL8⁻ |
| 10 | 16, 25 | CD3G⁺ |
| 11 | 14, 23 | TP63⁺ |

G

| Merged K-Means Clusters | Original K-Means Clusters | Marker |
|---|---|---|
| 1 | 2 | MAG⁺ |
| 2 | 3, 19 | CX3CR1⁺ |
| 3 | 7, 15 | CXCL8⁺ |
| 4 | 6 | DLL3⁺, MKI67⁺ |
| 5 | 9, 14 | TP73⁺ |
| 6 | 17 | EGFR⁺, MKI67⁺ |
| 7 | 8, 10, 13 | DLL3⁺, MKI67⁻ |
| 8 | 4, 16, 18 | EGFR⁺, MKI67⁻ |
| 9 | 12, 20 | CD44⁺, CXCL8⁻ |
| 10 | 1, 5 | CD3G⁺ |
| 11 | 11 | TP63⁺ |

**Supplementary Fig. S7.** Comparative analysis of dimension reduction analysis by Seurat based regular gene expression matrix and SCENIC based regulon matrix.

(A) SCENIC t-SNE colored with Seurat clusters; (B) K-Means clusters; (C) Merged Seurat clusters and (D) Merged K-Means clusters. Adjusted rand score and adjusted mutual info score are calculated and listed in two groups: (A) and (B), (C) and (D). (F-G) Table of merged Seurat and regulon K-Means clusters, based on the marker gene expression pattern showed in (E).

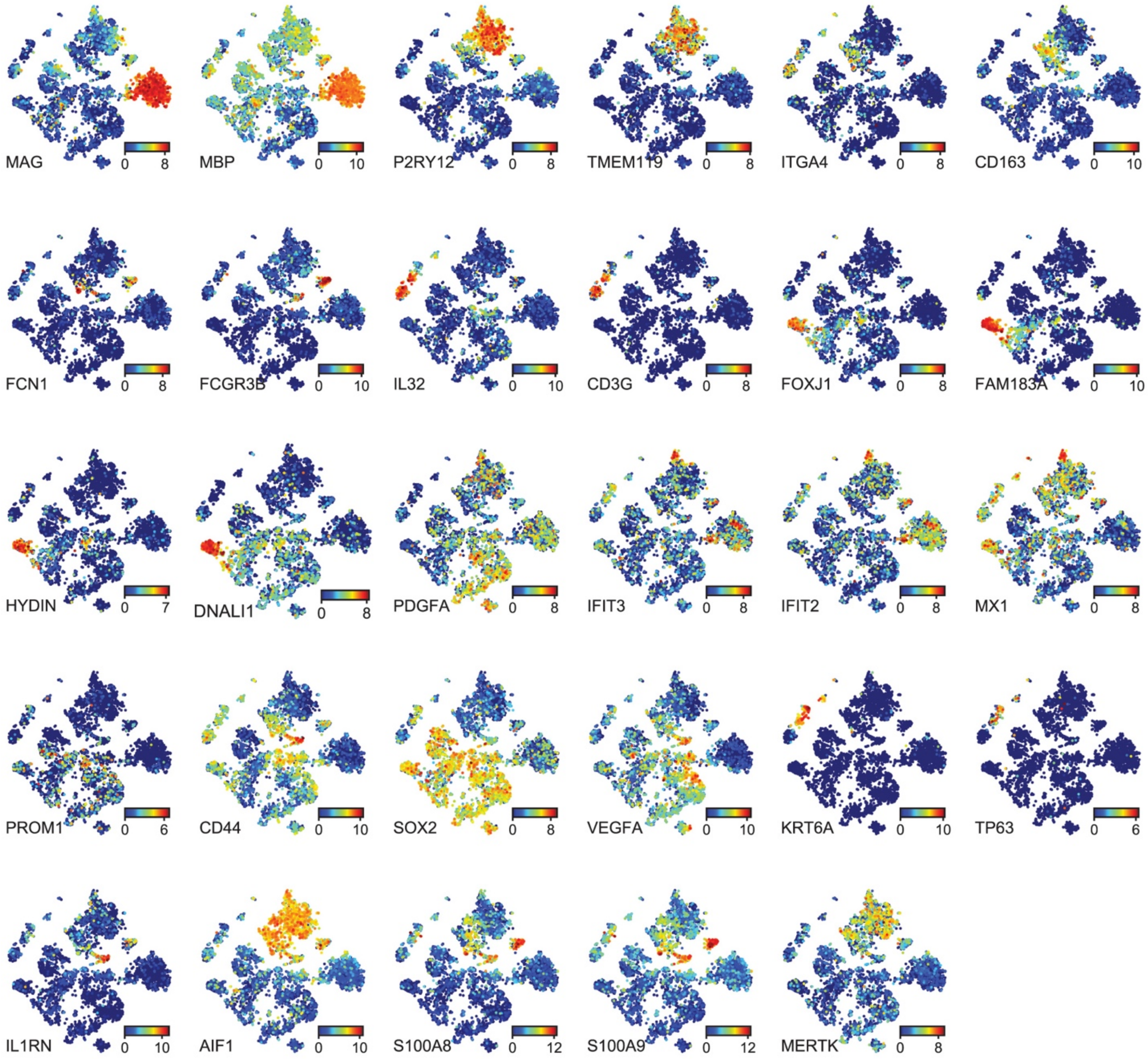
MAG 0 8
MBP 0 10
P2RY12 0 8
TMEM119 0 8
ITGA4 0 8
CD163 0 10
FCN1 0 8
FCGR3B 0 10
IL32 0 10
CD3G 0 8
FOXJ1 0 8
FAM183A 0 10
HYDIN 0 7
DNALI1 0 8
PDGFA 0 8
IFIT3 0 8
IFIT2 0 8
MX1 0 8
PROM1 0 6
CD44 0 10
SOX2 0 8
VEGFA 0 10
KRT6A 0 10
TP63 0 6
IL1RN 0 10
AIF1 0 10
S100A8 0 12
S100A9 0 12
MERTK 0 8

**Supplementary Fig. S8.** Marker gene expression feature map for cell type identification.

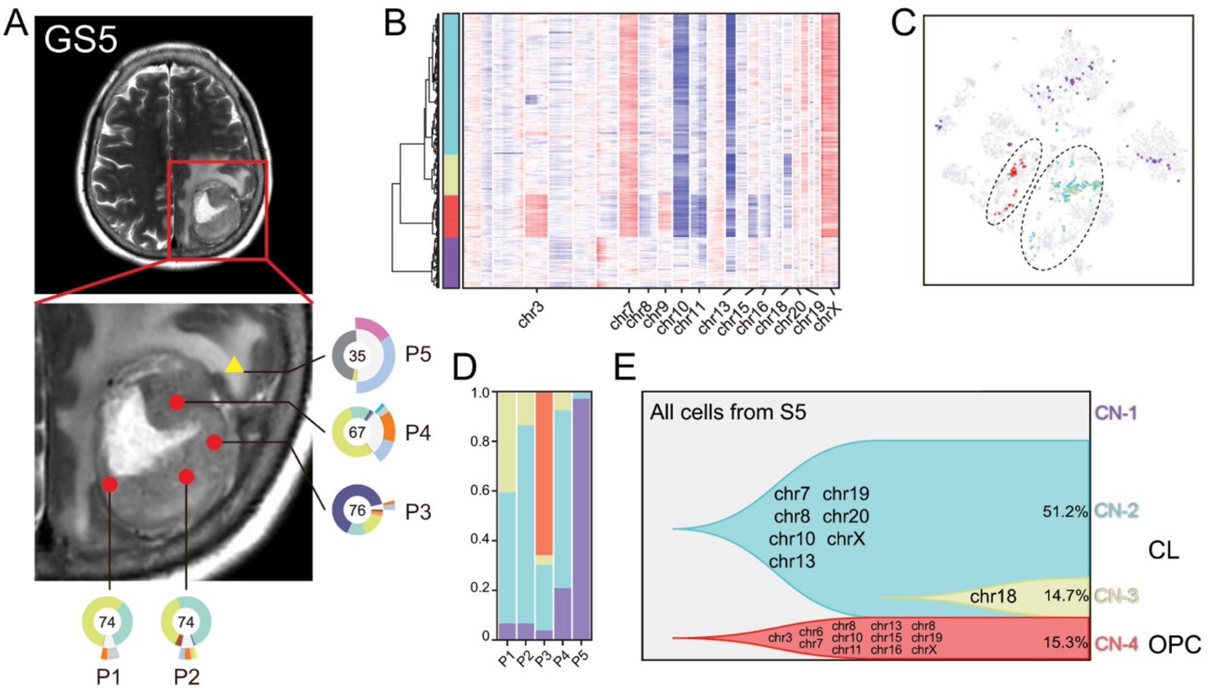

A
GS5
P5
35
P4
67
P3
76
74
74
P1
P2
B
chr3
chr7
chr8
chr9
chr10
chr11
chr13
chr15
chr16
chr18
chr20
chr19
chrX
C
D
1.0
0.8
0.6
0.4
0.2
0
P1
P2
P3
P4
P5
E
All cells from S5
chr7 chr19
chr8 chr20
chr10 chrX
chr13
chr18
chr3 chr6 chr7 chr8 chr10 chr11 chr13 chr15 chr16 chr8 chr19 chrX
51.2%
14.7%
15.3%
CN-1
CN-2
CN-3
CN-4
CL
OPC

**Supplementary Fig. S9.** Glioma clonal evolution of Patient GS5 at spatial and temporal resolution. (A) MRI image of Patient GS5. Yellow and red markers in zoomed image represent peritumoral and tumoral sampling points. Ring plot in the right and bottom displayed cell component of each point. Color of the inner ring showed classified glial cell subtypes, and the outer ring showed detailed immune cell subtypes. Cell numbers were labeled in the center of these ring plots. (B) Single-cell CNV heatmap. Cells were divided into 4 groups by hierarchical clustering. (C) CNV subclone distributions in t-SNE coordinates. (D) CNV subclone components in each sampling point. (E) Clonal evolution trail followed by accumulating CNV events. Each color represented a CNV subclone and chromosomes were labeled, in which copy number alteration were happened during clonal transition.

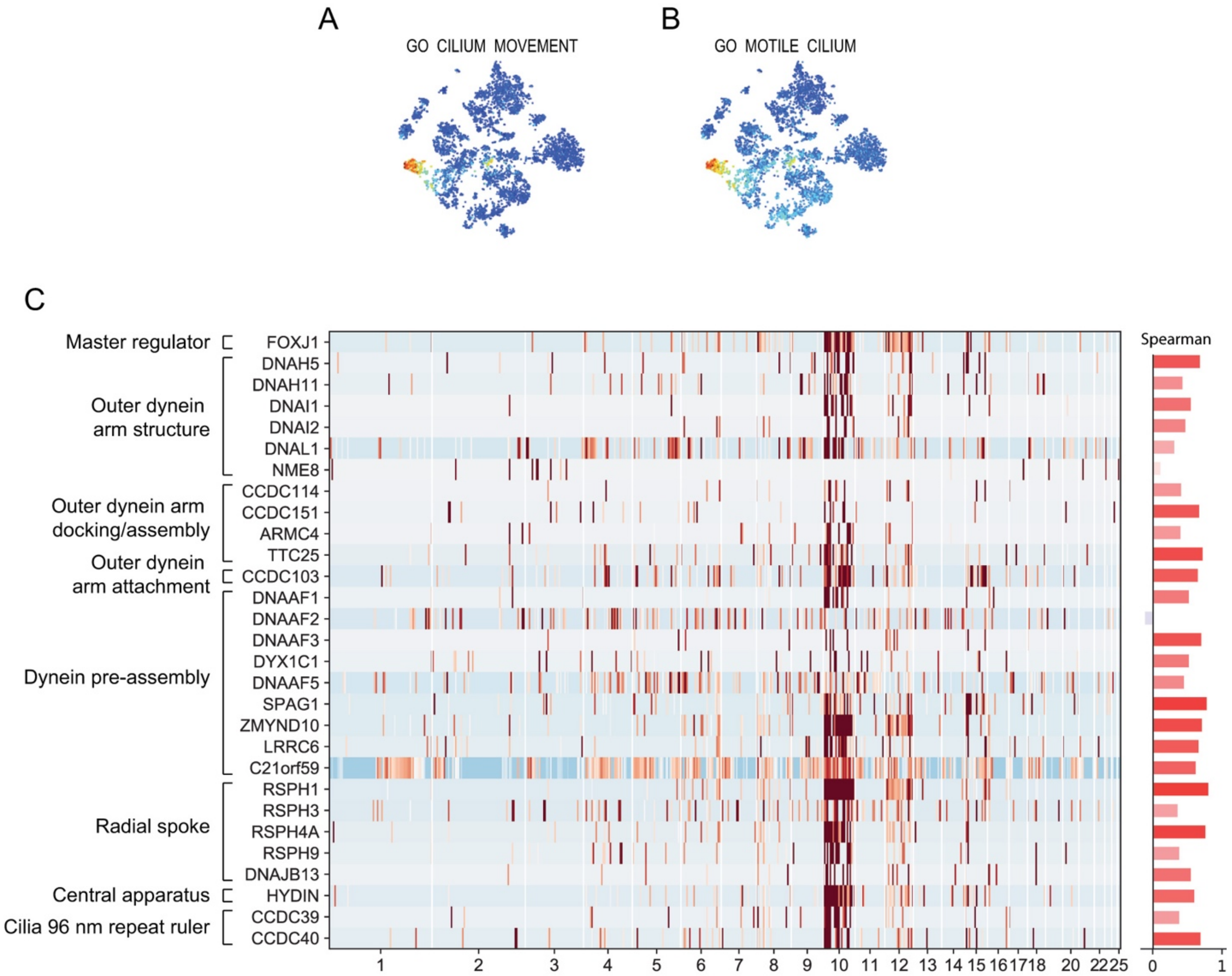
A
GO CILIUM MOVEMENT
B
GO MOTILE CILIUM
C
Master regulator
Outer dynein arm structure
Outer dynein arm docking/assembly
Outer dynein arm attachment
Dynein pre-assembly
Radial spoke
Central apparatus
Cilia 96 nm repeat ruler
FOXJ1
DNAH5
DNAH11
DNAI1
DNAI2
DNAL1
NME8
CCDC114
CCDC151
ARMC4
TTC25
CCDC103
DNAAF1
DNAAF2
DNAAF3
DYX1C1
DNAAF5
SPAG1
ZMYND10
LRRC6
C21orf59
RSPH1
RSPH3
RSPH4A
RSPH9
DNAJB13
HYDIN
CCDC39
CCDC40
1 2 3 4 5 6 7 8 9 10 11 12 13 14 15 16 17 18 20 22 25
Spearman
0 1

**Supplementary Fig. S10.** GS13 cilium cells overexpress motile cilium related marker genes. (A-B) Gene set score of motile cilium related GO terms. (C) Left panel: Expression heatmap of motile cilium related marker genes. Right panel: Spearman correlation coefficient values of marker genes to the master regulator *FOXJ1*.

A

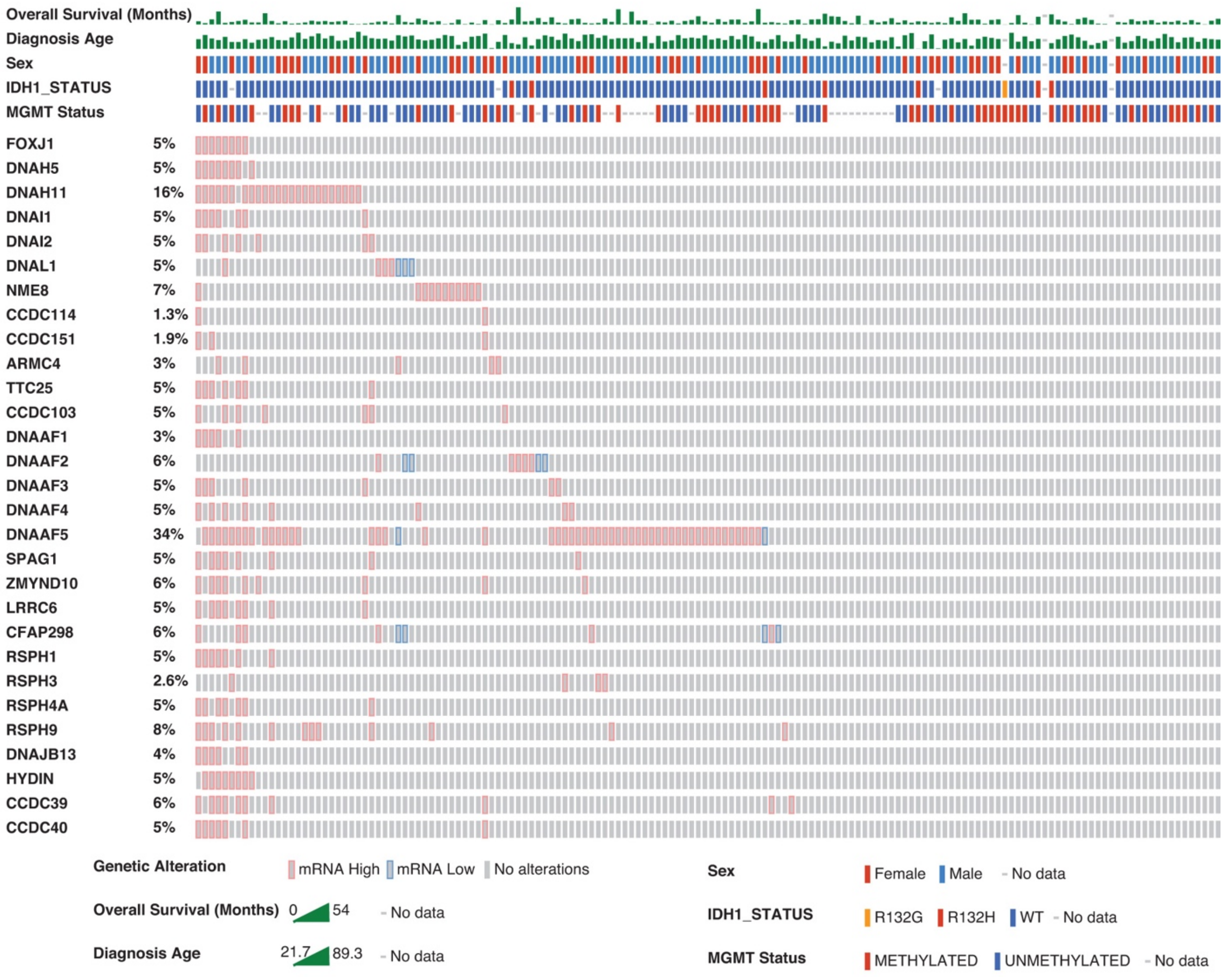
Overall Survival (Months)
Diagnosis Age
Sex
IDH1_STATUS
MGMT Status
FOXJ1 5%
DNAH5 5%
DNAH11 16%
DNAI1 5%
DNAI2 5%
DNAL1 5%
NME8 7%
CCDC114 1.3%
CCDC151 1.9%
ARMC4 3%
TTC25 5%
CCDC103 5%
DNAAF1 3%
DNAAF2 6%
DNAAF3 5%
DNAAF4 5%
DNAAF5 34%
SPAG1 5%
ZMYND10 6%
LRRC6 5%
CFAP298 6%
RSPH1 5%
RSPH3 2.6%
RSPH4A 5%
RSPH9 8%
DNAJB13 4%
HYDIN 5%
CCDC39 6%
CCDC40 5%
Genetic Alteration
mRNA High
mRNA Low
No alterations
Overall Survival (Months) 0 54 No data
Diagnosis Age 21.7 89.3 No data
Sex
Female
Male
No data
IDH1_STATUS
R132G
R132H
WT
No data
MGMT Status
METHYLATED
UNMETHYLATED
No data

B

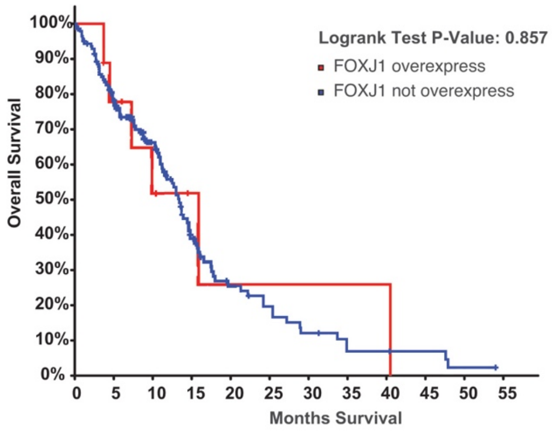
Logrank Test P-Value: 0.857
FOXJ1 overexpress
FOXJ1 not overexpress
Overall Survival
100%
90%
80%
70%
60%
50%
40%
30%
20%
10%
0%
0 5 10 15 20 25 30 35 40 45 50 55
Months Survival

**Supplementary Fig. S11.** Motile cilium marker overexpression samples in TCGA GBM cohort. (A) Overexpressed motile cilium related genes in TCGA GBM cohort. (B) Overall survival analysis of *FOXJ1* overexpressed TCGA GBM patients.

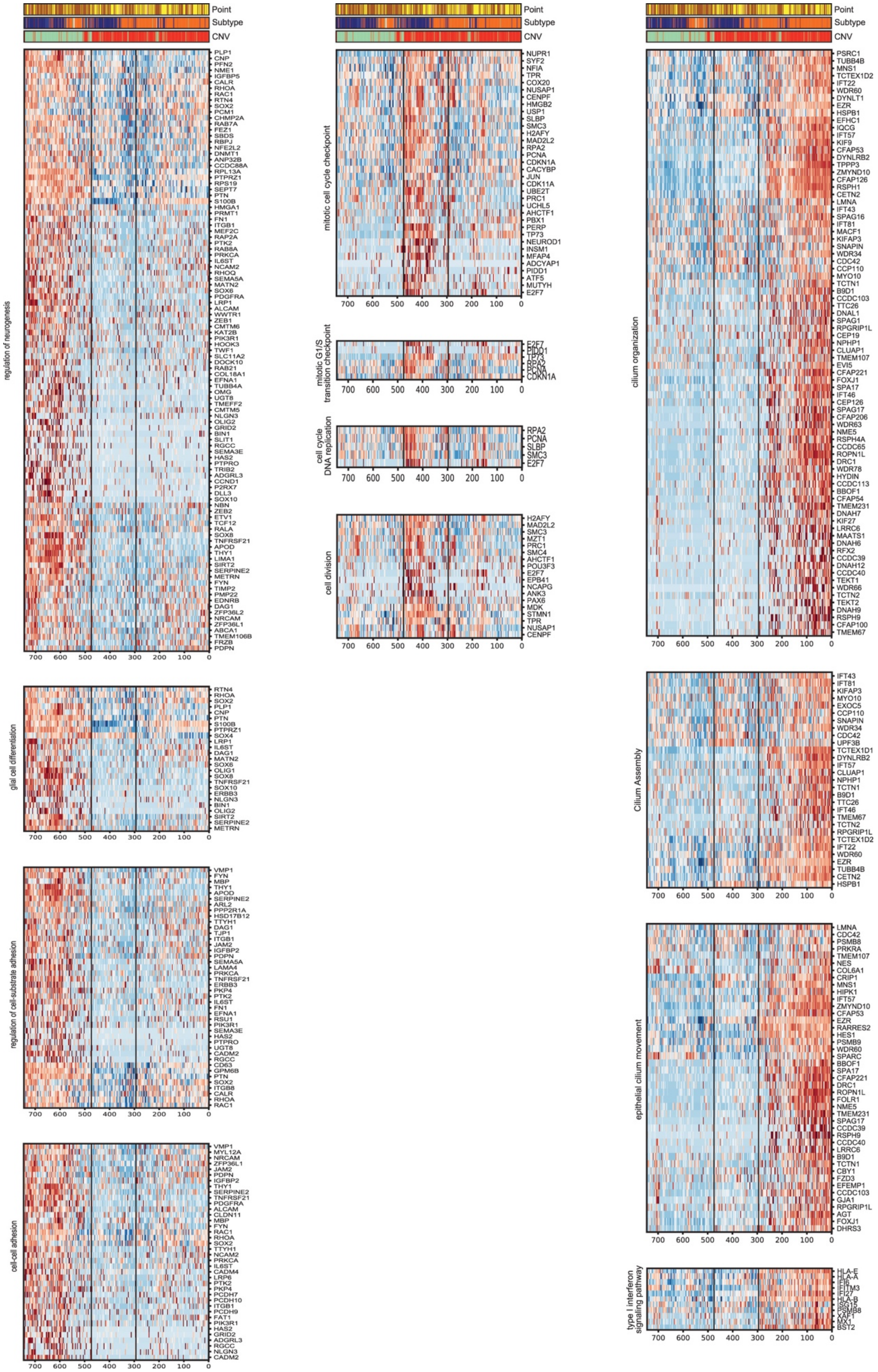

Point
Subtype
CNV
regulation of neurogenesis
PLP1
CNP
PFN2
NME1
IGFBP5
CALR
RHOA
RAC1
RTN4
SOX2
PCM1
CHMP2A
RAB7A
FEZ1
SBDS
RBPJ
NFE2L2
DNMT1
ANP32B
CCDC88A
RPL13A
PTPRZ1
RPS19
SEPT7
PTN
S100B
HMGA1
PRMT1
FN1
ITGB1
MEF2C
RAP2A
PTK2
RAB8A
PRKCA
IL6ST
NCAM2
RHOQ
SEMA5A
MATN2
SOX6
PDGFRA
LRP1
ALCAM
WWTR1
ZEB1
CMTM6
KAT2B
PIK3R1
HOOK3
TWF1
SLC11A2
DOCK10
RAB21
COL18A1
EFNA1
TUBB4A
OMG
UGT8
TMEFF2
CMTM5
NLGN3
OLIG2
GRID2
BIN1
SLIT1
RGCC
SEMA3E
HAS2
PTPRO
TRIB2
ADGRL3
CCND1
P2RX7
DLL3
SOX10
NBN
ZEB2
ETV1
TCF12
RALA
SOX8
TNFRSF21
APOD
THY1
LIMA1
SIRT2
SERPINE2
METRN
FYN
TIMP2
PMP22
EDNRB
DAG1
ZFP36L2
NRCAM
ZFP36L1
ABCA1
TMEM106B
FRZB
PDPN
700 600 500 400 300 200 100 0
glial cell differentiation
RTN4
RHOA
SOX2
PLP1
CNP
PTN
S100B
PTPRZ1
SOX4
LRP1
IL6ST
DAG1
MATN2
SOX6
OLIG1
SOX8
TNFRSF21
SOX10
ERBB3
NLGN3
BIN1
OLIG2
SIRT2
SERPINE2
METRN
700 600 500 400 300 200 100 0
regulation of cell-substrate adhesion
VMP1
FYN
MBP
THY1
APOD
SERPINE2
ARL2
PPP2R1A
HSD17B12
TTYH1
DAG1
TJP1
ITGB1
JAM2
IGFBP2
PDPN
SEMA5A
LAMA4
PRKCA
TNFRSF21
ERBB3
PKP4
PTK2
IL6ST
FN1
EFNA1
RSU1
PIK3R1
SEMA3E
HAS2
PTPRO
UGT8
CADM2
RGCC
CD63
GPM6B
PTN
SOX2
ITGB8
CALR
RHOA
RAC1
700 600 500 400 300 200 100 0
cell-cell adhesion
VMP1
MYL12A
NRCAM
ZFP36L1
JAM2
PDPN
IGFBP2
THY1
SERPINE2
TNFRSF21
PDGFRA
ALCAM
CLDN11
MBP
FYN
RAC1
RHOA
SOX2
TTYH1
NCAM2
PRKCA
IL6ST
CADM4
LRP6
PTK2
PKP4
PCDH7
PCDH10
ITGB1
PCDH9
FAT1
PIK3R1
HAS2
GRID2
ADGRL3
RGCC
NLGN3
CADM2
700 600 500 400 300 200 100 0
mitotic cell cycle checkpoint
NUPR1
SYF2
NFIA
TPR
COX20
NUSAP1
CENPF
HMGB2
USP1
SLBP
SMC3
H2AFY
MAD2L2
RPA2
PCNA
CDKN1A
CACYBP
JUN
CDK11A
UBE2T
PRC1
UCHL5
AHCTF1
PBX1
PERP
TP73
NEUROD1
INSM1
MFAP4
ADCYAP1
PIDD1
ATF5
MUTYH
E2F7
700 600 500 400 300 200 100 0
mitotic G1/S transition checkpoint
E2F7
PIDD1
TP73
RPA2
PCNA
CDKN1A
700 600 500 400 300 200 100 0
cell cycle DNA replication
RPA2
PCNA
SLBP
SMC3
E2F7
700 600 500 400 300 200 100 0
cell division
H2AFY
MAD2L2
SMC3
MZT1
PRC1
SMC4
AHCTF1
POU3F3
E2F7
EPB41
NCAPG
ANK3
PAX6
MDK
STMN1
TPR
NUSAP1
CENPF
700 600 500 400 300 200 100 0
cilium organization
PSRC1
TUBB4B
MNS1
TCTEX1D2
IFT22
WDR60
DYNLT1
EZR
HSPB1
EFHC1
IQCG
IFT57
KIF9
CFAP53
DYNLRB2
TPPP3
ZMYND10
CFAP126
RSPH1
CETN2
LMNA
IFT43
SPAG16
IFT81
MACF1
KIFAP3
SNAPIN
WDR34
CDC42
CCP110
MYO10
TCTN1
B9D1
CCDC103
TTC26
DNAL1
SPAG1
RPGRIP1L
CEP19
NPHP1
CLUAP1
TMEM107
EVI5
CFAP221
FOXJ1
SPA17
IFT46
CEP126
SPAG17
CFAP206
WDR63
NME5
RSPH4A
CCDC65
ROPN1L
DRC1
WDR78
HYDIN
CCDC113
BBOF1
CFAP54
TMEM231
DNAH7
KIF27
LRRC6
MAATS1
DNAH6
RFX2
CCDC39
DNAH12
CCDC40
TEKT1
WDR66
TCTN2
TEKT2
DNAH9
RSPH9
CFAP100
TMEM67
700 600 500 400 300 200 100 0
Cilium Assembly
IFT43
IFT81
KIFAP3
MYO10
EXOC5
CCP110
SNAPIN
WDR34
CDC42
UPF3B
TCTEX1D1
DYNLRB2
IFT57
CLUAP1
NPHP1
TCTN1
B9D1
TTC26
IFT46
TMEM67
TCTN2
RPGRIP1L
TCTEX1D2
IFT22
WDR60
EZR
TUBB4B
CETN2
HSPB1
700 600 500 400 300 200 100 0
epithelial cilium movement
LMNA
CDC42
PSMB8
PRKRA
TMEM107
NES
COL6A1
CRIP1
MNS1
HIPK1
IFT57
ZMYND10
CFAP53
EZR
RARRES2
HES1
PSMB9
WDR60
SPARC
BBOF1
SPA17
CFAP221
DRC1
ROPN1L
FOLR1
NME5
TMEM231
SPAG17
CCDC39
RSPH9
CCDC40
LRRC6
B9D1
TCTN1
CBY1
FZD3
EFEMP1
CCDC103
GJA1
RPGRIP1L
AGT
FOXJ1
DHRS3
700 600 500 400 300 200 100 0
type I interferon signaling pathway
HLA-E
HLA-A
IFI6
IFITM3
IFI27
HLA-B
ISG15
PSMB8
XAF1
MX1
BST2
700 600 500 400 300 200 100 0

**Supplementary Fig. S12.** Gene expression heatmap of individual genes comprised in the GO terms listed in Figure 5G.

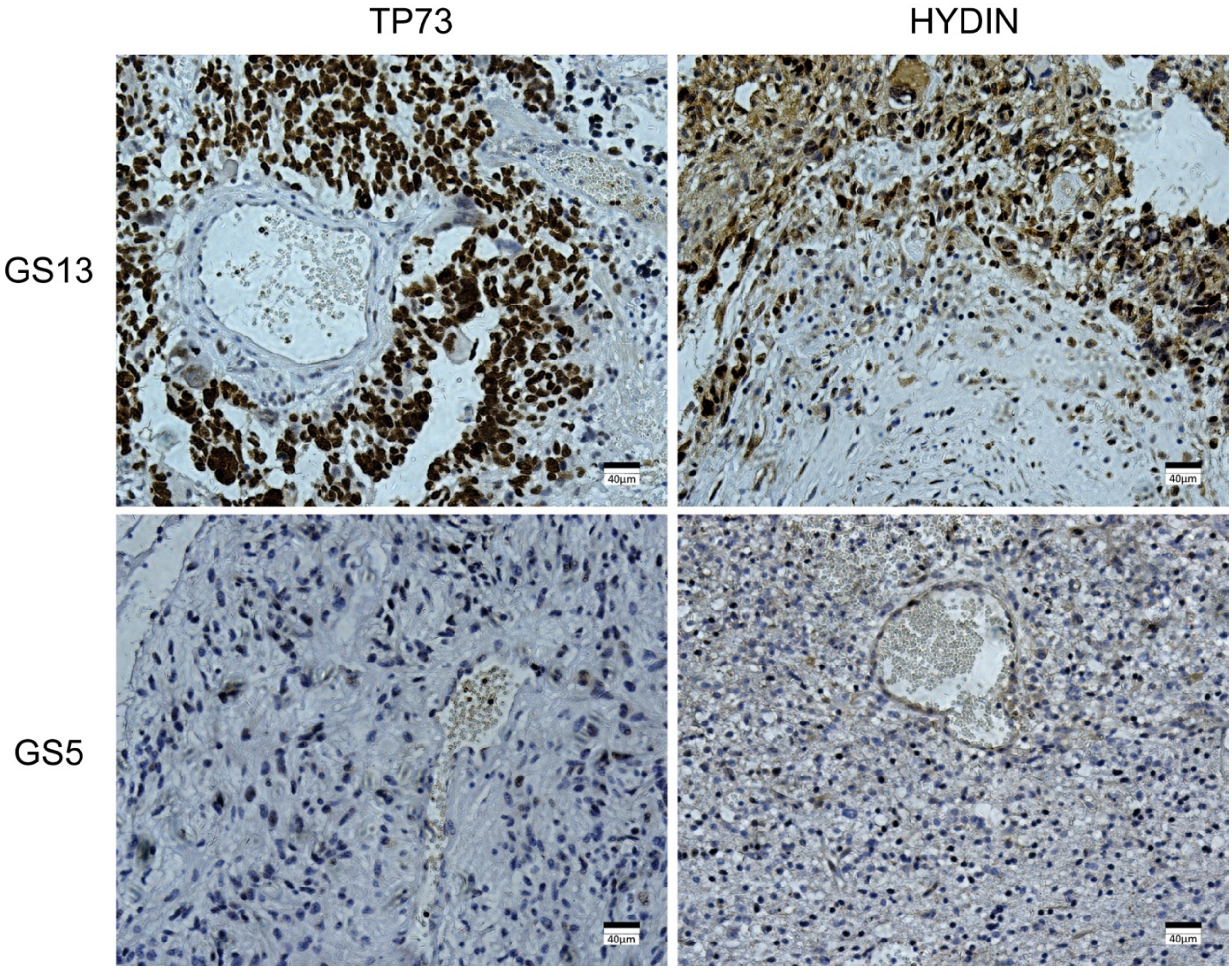
TP73
HYDIN
GS13
GS5
40μm
40μm
40μm
40μm

**Supplementary Fig. S13.** Immunohistochemistry staining image of *TP73* and *HYDIN* protein on paraffin slices of GS13 and GS5.

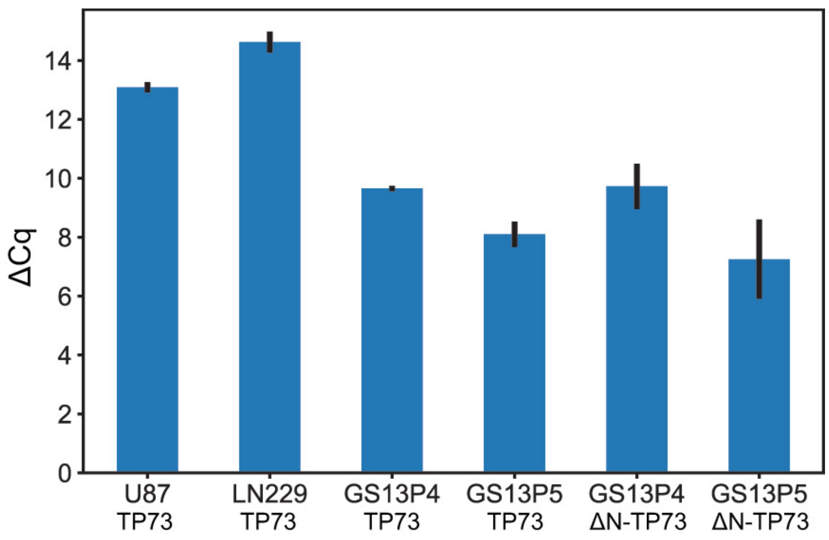

ΔCq
0
2
4
6
8
10
12
14
U87 TP73
LN229 TP73
GS13P4 TP73
GS13P5 TP73
GS13P4 ΔN-TP73
GS13P5 ΔN-TP73

**Supplementary Fig. S14.** Expression level of full length *TP73* and *ΔN-TP73* isoform. RT-qPCR of full length *TP73* and *ΔN-TP73* isoform. Both of them exists in GS13 cilium cells.

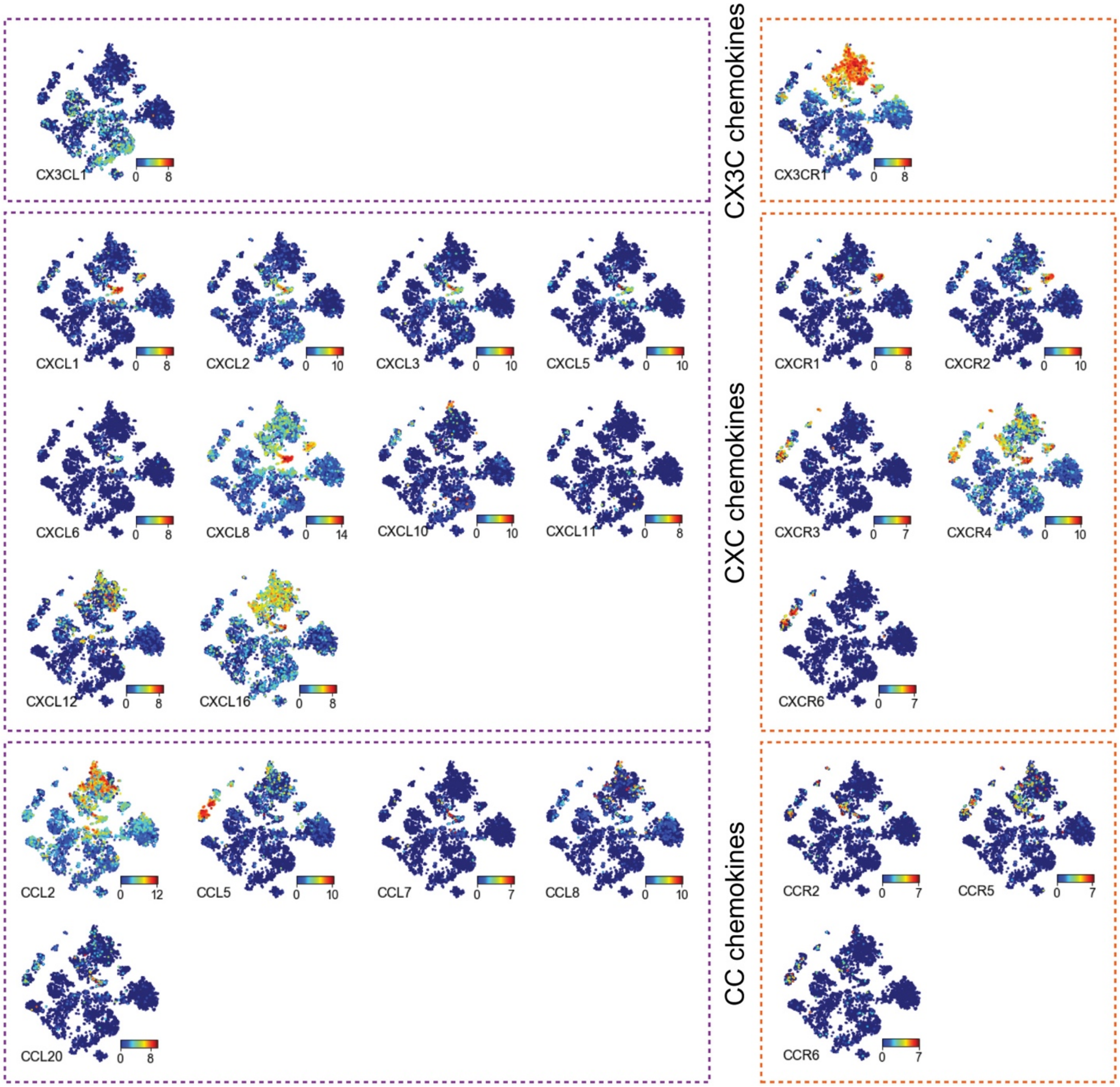
CX3C chemokines
CXC chemokines
CC chemokines
CX3CL1
CX3CR1
CXCL1
CXCL2
CXCL3
CXCL5
CXCL6
CXCL8
CXCL10
CXCL11
CXCL12
CXCL16
CXCR1
CXCR2
CXCR3
CXCR4
CXCR6
CCL2
CCL5
CCL7
CCL8
CCL20
CCR2
CCR5
CCR6

**Supplementary Fig. S15.** Cell source information of each patient in the pseudotime map from Fig. 6A**.**

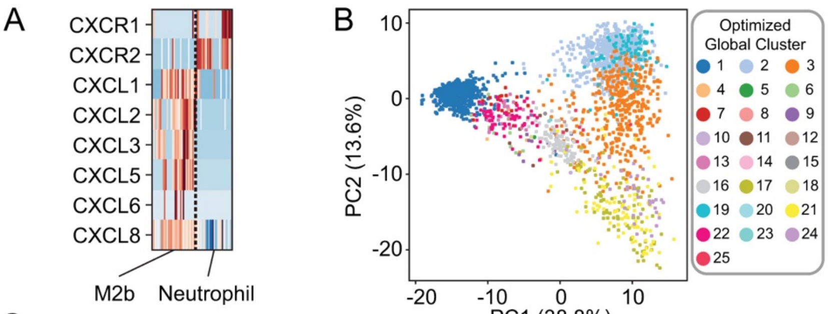


C

| | | | | | | | | | |
|---|---|---|---|---|---|---|---|---|---|
| AQP9 | BCL2A1 | C15orf48 | C5AR1 | CD55 | CEBPB | CXCL1 | CXCR2 | FCGR3B | FCN1 |
| FGR | FNDC3B | IFITM2 | IL1R2 | IL1RN | LILRB2 | LILRB3 | LITAF | NAMPT | NCF1C |
| NCF2 | PLAUR | RNF149 | S100A12 | S100A8 | S100A9 | SDCBP | SERPINA1 | SLC2A3 | SOCS3 |
| SOD2 | SPI1 | TCIRG1 | THBD | TREM1 | UPP1 | VASP | VNN2 | | |

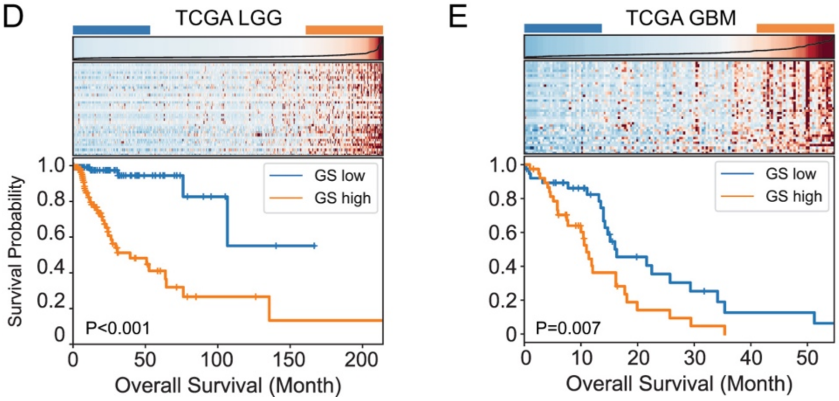

**Supplementary Fig. S16.** Characteristics of m2b macrophages and neutrophils and their potential in prognosis prediction using M2bNeu score

(A) Expression profile of chemotaxis related genes in M2b and neutrophil cells. (B) First 2 PCs of TME cell PCA analysis, using the same color code as (Fig. 2D) cell type color bar. M2b and neutrophils have similar characteristics in PC2. (C) M2bNeu gene list. (D-E) Prognosis prediction using the M2bNeu score. The scores were calculated with TCGA LGG/GBM bulk RNA-seq data. For each of these two groups of patients, samples in the upper quantiles (orange) of the ranked M2bNeu score showed a significantly worse prognosis than those in the lower quantiles (blue).

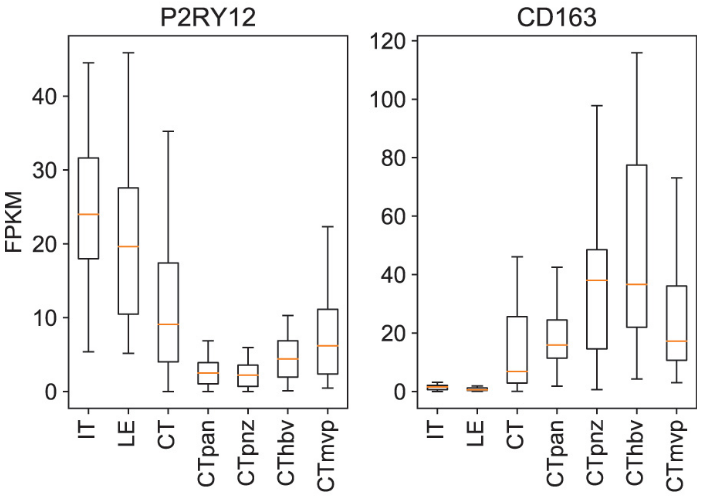

P2RY12
CD163
FPKM
0
10
20
30
40
0
20
40
60
80
100
120
IT
LE
CT
CTpan
CTpnz
CThbv
CTmvp
IT
LE
CT
CTpan
CTpnz
CThbv
CTmvp

**Supplementary Fig. S17.** *P2RY12* and *CD163* expression in IVY GAP database.
5 major anatomic structures of glioblastoma: Leading Edge (LE), Infiltrating Tumor (IT), Cellular Tumor (CT), Pseudopalisading Cells Around Necrosis (pan), Peri-Necrotic Zone (pnz), Hyperplastic Blood Vessels (hbv), Microvascular Proliferation (mvp).

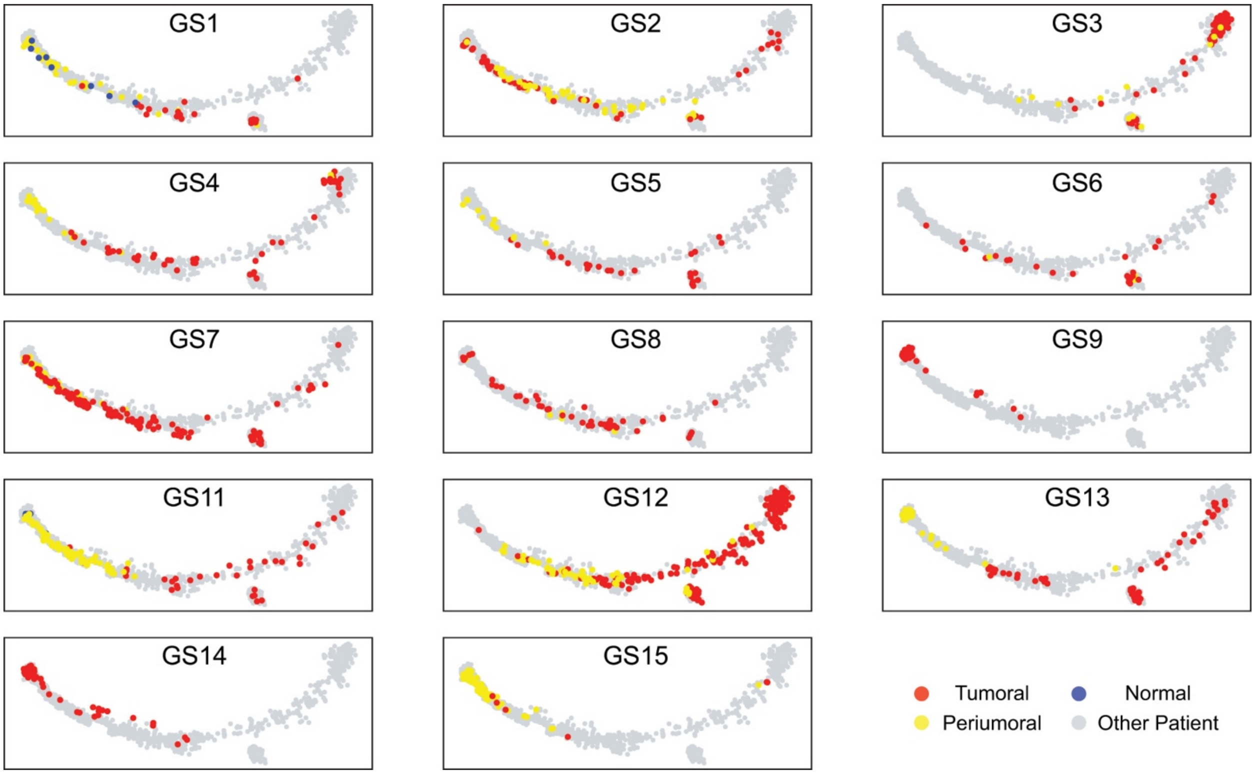

GS1
GS2
GS3
GS4
GS5
GS6
GS7
GS8
GS9
GS11
GS12
GS13
GS14
GS15
Tumoral
Normal
Periumoral
Other Patient

**Supplementary Fig. S18.** Cell source information of each patient in the pseudotime map from Fig. 6D.

#### Supplementary Tables

**Supplementary Table 1.** Clinical information.
Clinical information of 14 patients and information of all biopsies.

**Supplementary Table 2.** Gene lists.
Marker genes of 25 clusters in the global t-SNE analysis and clustered marker genes related to Fig. 4F, 5F.
